# CLSTN3B enhances adipocyte lipid droplet structure and function via endoplasmic reticulum contact

**DOI:** 10.1101/2024.01.20.576491

**Authors:** Chuanhai Zhang, Mengchen Ye, Kamran Melikov, Dengbao Yang, Goncalo Dias do Vale, Jeffrey McDonald, Kaitlyn Eckert, Mei-Jung Lin, Xing Zeng

## Abstract

Interorganelle contacts facilitate material exchanges and sustain the structural and functional integrity of organelles. Lipid droplets (LDs) of adipocytes are responsible for energy storage and mobilization responding to body needs. LD biogenesis defects compromise the lipid-storing capacity of adipocytes, resulting in ectopic lipid deposition and metabolic disorders, yet how the uniquely large LDs in adipocytes attain structural and functional maturation is incompletely understood. Here we show that the mammalian adipocyte-specific protein CLSTN3B is crucial for adipocyte LD maturation. CLSTN3B employs an arginine-rich segment to promote extensive contact and hemifusion-like structure formation between the endoplasmic reticulum (ER) and LD, allowing ER-to-LD phospholipid diffusion during LD expansion. CLSTN3B ablation results in reduced LD surface phospholipid density, increased turnover of LD-surface proteins, and impaired LD functions. Our results establish the central role of CLSTN3B in the adipocyte-specific LD maturation pathway that enhances lipid storage and maintenance of metabolic health under caloric overload.

---

Adipocytes store excess energy in the form of neutral lipids, primarily triacylglycerides (TG). Insufficient lipid-storing capacity of adipocytes results in ectopic lipid deposition in extra-adipose organs and drives the pathogenesis of metabolic disorders, such as type II diabetes^1^. Enhancing lipid storage in adipocytes with peroxisome proliferator activated receptor gamma (PPARG) agonists improves insulin sensitivity^2,3^. However, the clinical use of PPARG agonists has been hampered by cardiovascular side effects^4^, calling for the elucidation of additional limiting factors governing the lipid-storing capacity of adipocytes.

Cells store neutral lipids in lipid droplets (LDs), organelles coated with a monolayer of phospholipids and various LD-targeting proteins. LD surface structures protect the internal lipid storage from unregulated lipolysis while sensitizing lipid mobilization to hormonal stimulation^5,6^. LD biogenesis occurs at the endoplasmic reticulum (ER) and ER-LD contact sites mediate protein and lipid transfer from the ER to LDs^7–15^. Additional LD-targeting proteins are directly recruited from the cytosol^16^. Impaired LD biogenesis or functionality compromises the lipid-storage capacity of adipocytes and increases susceptibility to metabolic diseases^17–19^. How adipocytes employ cell type-specific mechanisms to attain structural and functional maturation of their uniquely large and dynamic LDs remains to be fully understood.

Here we show that the mammalian adipocyte-specific protein Calsyntenin 3β (CLSTN3B) works at the ER/LD interface to promote LD maturation. We found that CLSTN3B facilitates ER-to-LD phospholipid diffusion during LD expansion via extensive ER/LD contact and hemifusion-like structure. CLSTN3B deficiency results in reduced LD surface phospholipid density and increased turnover of LD-targeting proteins. Consequently, ablating CLSTN3B from white adipocytes results in reduced lipid-storing capacity of the white adipose tissue (WAT) and increases susceptibility to ectopic lipid deposition and onset of metabolic disorders at the animal level, whereas enhancing CLSTN3B expression in white adipocytes yields the opposite phenotype. Our results demonstrate how CLSTN3B-mediated inter-organelle communication promotes LD maturation, enhances lipid storage in adipocytes, and translates into physiological significance at the animal level.

## CLSTN3B enhances lipid storage in adipocytes

We previously showed that CLSTN3B is highly expressed in brown adipocytes and contributes to thermogenesis^20^. To probe the physiological relevance of CLSTN3B to white adipocytes primarily responsible for TG storage but not thermogenesis, we employed a condition favoring TG deposition: high-fat diet (HFD) feeding at thermoneutrality (28-30C). Under this condition, *clstn3b^-/-^*mice had significantly reduced perigonadal white adipose tissue (pgWAT) masses, smaller adipocyte sizes in the pgWAT, more severe liver steatosis, higher serum free fatty acid (FFA) levels, and concomitant glucose intolerance compared with WT mice (Fig. 1a-g). Similar phenotypes were observed in both sexes and at room temperature and unlikely resulted from defective adipogenesis and lipodystrophy (Extended Data Fig. 1 and 2). The reduced pgWAT mass is unlikely caused by defective sympathetic innervation as previously noted in the BAT of the *clstn3b^-/-^* mice^20^, because innervation deficiency is expected to cause increased lipid accumulation, contrary to the reduced pgWAT masses as observed. Furthermore, pgWAT receives minimal parenchymal sympathetic innervation, which is not affected by CLSTN3B ablation (Extended Data Fig. 3). Restoring CLSTN3B expression specifically in brown adipocytes of the *clstn3b^-/-^* mice as previously published failed to improve the metabolic parameters^20^ (Extended Data Fig. 4a-d), indicating that CLSTN3B deficiency in white adipocytes is primarily responsible for the phenotype. As an alternative approach to delineate the contribution from brown and white adipocyte-derived CLSTN3B to the metabolic phenotypes, we constructed pan-adipocyte CLSTN3B and brown adipocyte-specific CLSTN3B knockout (KO) mice (*adq-cre*, *clstn3b^fl/fl^*; *ucp1-cre*, *clstn3b^fl/fl^*) and subjected them to HFD feeding at thermoneutrality. Whereas the pan-adipocyte CLSTN3B KO mice displayed similar metabolic defects to the global *clstn3b^-/-^* mice, the brown adipocyte-specific KO mice were indistinguishable from their WT control littermates (Extended Data Fig. 4e-n). These data again support that CLSTN3B expression in white adipocytes are responsible for enhancing lipid storage in the WAT.

**Figure 1.**
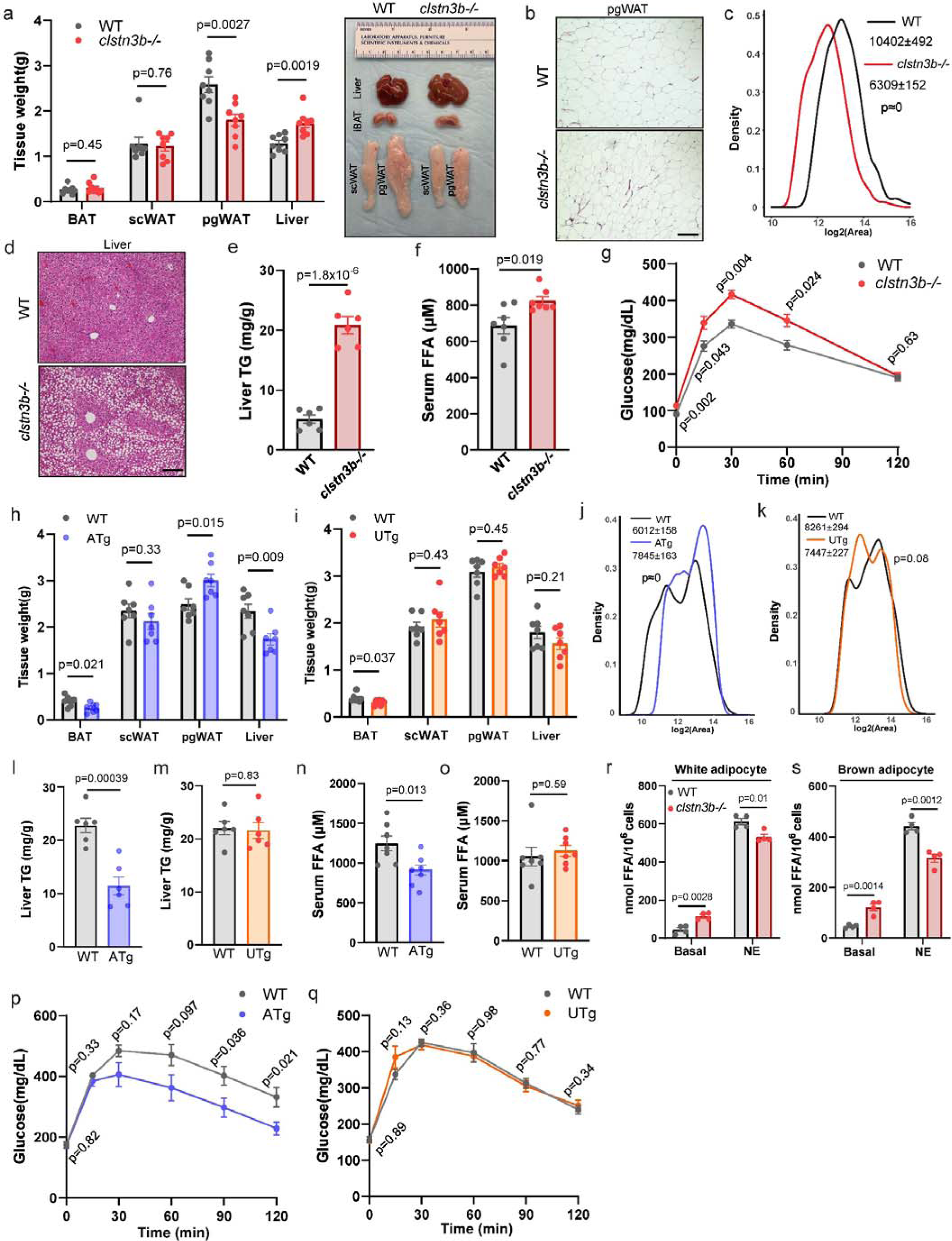
CLSTN3B enhances lipid storage in adipocytes. **a-g**, Tissue weights and gross appearances (**a**), pgWAT histology (**b**), pgWAT adipocyte size distributions (**c**), liver histology (**d**), liver TG contents (**e**), serum FFA (**f**), and glucose tolerance test (**g**) of WT and *clstn3b^-/-^* mice on HFD at thermoneutrality (n=7 mice). **h**-**q**, Tissue weights (**h, i**), pgWAT adipocyte size distributions (**j, k**), liver TG contents (**l, m**), serum FFA (**n, o**), and glucose tolerance test (**p, q**) of *adq-cre clstn3b* transgenic mice, *ucp1-cre clstn3b* transgenic mice and WT littermates on HFD at thermoneutrality (n=7 mice). **r-s**, Lipolysis assay of WT and *clstn3b^-/-^* white (**r**) and brown (**s**) adipocytes (n=4 replicates). Scale bar: 100 μm.

To complement the loss-of-function models, we analyzed transgenic mice overexpressing CLSTN3B in all adipocytes (ATg, *adq-cre*, *clstn3b^tg/^*^0^) or only brown adipocytes (UTg, *ucp1-cre*, *clstn3b^tg/^*^0^) fed with HFD at thermoneutrality. Only the ATg but not the UTg mice displayed improved metabolic parameters compared with littermate control mice (Fig. 1h-q, Extended Data Fig. 5), again highlighting the requirement of CLSTN3B expression in white adipocytes for improving lipid-storage in the WAT and metabolic health.

We then performed lipolytic assays with isolated *clstn3b^-/-^* white adipocytes to probe the reason underlying reduced lipid-storage capacity and found that those cells exhibited higher basal lipolysis rates and blunted responses to norepinephrine (NE) stimulation with similar levels of Hormone Sensitive Lipase (HSL) phosphorylation to WT cells (Fig. 1r and Extended Data Fig. 6). These data suggest that *clstn3b^-/-^* white adipocytes have impaired LD functionality. *Clstn3b^-/-^* brown adipocytes displayed a similar phenotype (Fig. 1s), indicating that CLSTN3B may enhance LD functionality in all adipocytes.

## CLSTN3B enhances LD phospholipid and protein levels

To understand the molecular basis of the abnormal lipolytic behavior of *clstn3b^-/-^* adipocytes, we conducted a systematic analysis of the LD surface structure. We first measured LD surface phospholipid density (Extended Data Fig. 7) and found that the area per phospholipid molecule is 71.7 ± 3.5 Å^2^ on WT brown adipocyte LD and 88.3 ± 4.6 Å^2^ on KO brown adipocyte LD (Fig. 2a). Similarly, the area per phospholipid molecule on WT white adipocyte LD is 79.8 ± 4.1 Å^2^ and 99.8 ± 4.2 Å^2^ on KO white adipocyte LD (Fig. 2a). Phospholipidomics analysis showed a uniform decrease across a wide range of phospholipid species (Extended Data Fig. 8). Taken together, these results show CLSTN3B ablation causes reduced phospholipid densities on adipocyte LDs.

**Figure 2.**
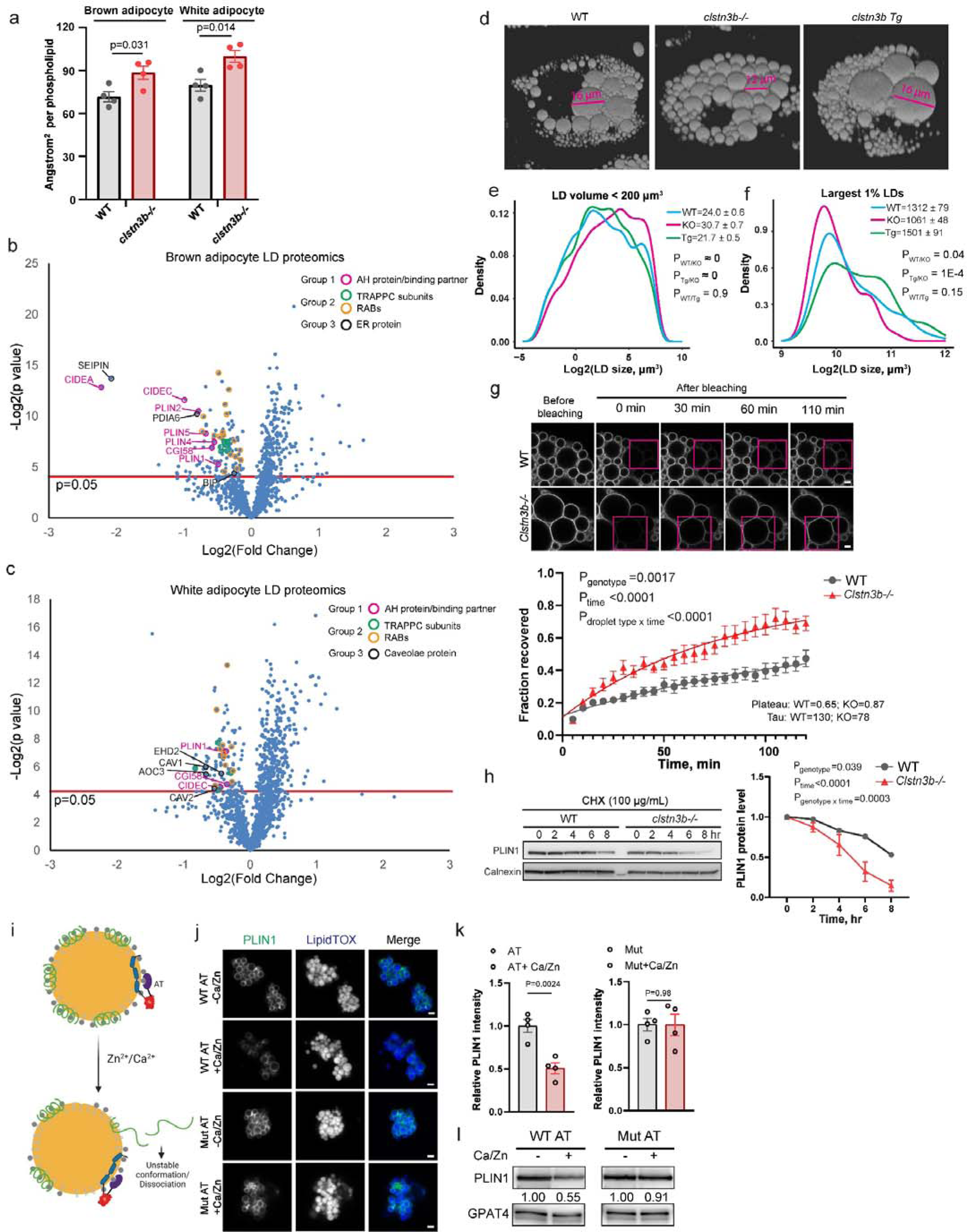
CLSTN3B enhances LD phospholipid and protein levels. **a**, Quantitation of area per phospholipid molecule on brown and white adipocyte LD isolated from WT and *clstn3b^-/-^* mice (n=4 replicates). **b-c**, Proteomics analysis of brown (**b**) and white (**c**) adipocyte LDs isolated from WT and *clstn3b^-/-^*mice (n=4 replicates). **d-f**, 3D reconstruction (**d**), size distribution of small (**e**) and large (**f**) LDs in WT, *clstn3b^-/-^* and transgenic brown adipocytes differentiated *in vitro*. **g**, FRAP analysis of PLIN1-mcherry turnover rate in WT and clstn3b-/- brown adipocytes. **h**, Measurement of PLIN1 half-life in WT and clstn3b-/- brown adipocytes by cycloheximide chase (n=3 replicates). **i-l**, Schematics (**i**), fluorescence image (**j**) and quantitation (**k**), and western blot (**l**) analysis of PLIN1-mclover3 levels on LDs with LD-targeted WT or catalytically inactive alpha-toxin -/+ Ca^2+^/Zn^2+^ activation. Scale bar: 5 μm.

We next performed proteomics analyses of adipocyte LDs isolated from WT and *clstn3b^-/-^* mice (Table S1). Significantly downregulated proteins in the *clstn3b^-/-^* brown adipocyte LD samples relative to the WT (Fig. 2b and Extended Data Fig. 9) included the following groups: 1) Amphipathic-helix (AH) containing LD-targeting proteins and their binding partners^16^; 2) Proteins involved in membrane fusion: including multiple RABs and associated Guanine Exchange Factors (GEFs); 3) ER integral membrane and lumenal proteins (BSCL2, PDIA6 and HSPA5). Proteomics analyses of white adipocyte LDs revealed similar downregulation of group 1 and 2 proteins but not group 3 (Fig. 2c and Extended Data Fig. 9). Interestingly, caveolae proteins were significantly downregulated in the *clstn3b^-/-^* white adipocyte LD samples (Fig. 2c), consistent with a previous report on the association of caveolae proteins with white adipocyte LDs^21^. The downregulation of PLIN1 was verified by Western blot (Extended Data Fig. 10) and may be particularly relevant to the abnormal lipolytic behavior of *clstn3b^-/-^* adipocytes and the metabolic phenotype of *clstn3b^-/-^* mice, because *plin1^-/-^* adipocytes and mice phenocopy their *clstn3b^-/-^* counterparts^22,23^. The mRNA levels of downregulated LD proteins were not significantly different between WT and *clstn3b^-/-^* brown or white adipocytes (Extended Data Fig. 11).

Decreased LD surface coverage by phospholipids increases LD surface tension and promotes LD coalescence^24^. Indeed, LDs below 200 μm^3^ in *clstn3b^-/-^* cells are larger than those in WT and Tg cells, consistent with increased coalescence of small LDs. However, the largest LDs in WT and Tg are larger than those in *clstn3b^-/-^* cells (Fig. 2, d-f). The largest LDs in brown adipocytes are formed via CIDE-protein facilitated assimilation of small LDs^25^. The significant downregulation of CIDEA and CIDEC on LDs from *clstn3b^-/-^* adipocytes (Fig. 2b-c) thus explains why the largest LDs in *clstn3b^-/-^* cells fail to become as large as in WT or Tg cells. (Fig. 2f). The simultaneous downregulation of phospholipids and AH containing proteins raises a conundrum because LDs with decreased phospholipid packing are expected to recruit more AH-containing proteins^26^. Previous studies demonstrated cooperative binding of AHs and phospholipids at the oil-water interface^27,28^. We therefore hypothesized that LD surface phospholipids may promote better retaining and reduced turnover of AH-containing proteins. This idea is supported by a shorter half-life of PLIN1 in *clstn3b^-/-^* brown adipocytes, a higher turnover rate of LD-bound PLIN1-mCherry in *clstn3b^-/-^* brown adipocyte, and a stronger propensity for a PLIN1 AH peptide to dissociate from neat triolein droplets than phospholipid-coated triolein droplets (Fig. 2g-h and Extended Data Fig. 12). Artificial reduction of LD surface phospholipid with an LD-targeting bacterial phospholipase C, also known as Alpha-Toxin, from *Clostridium perfringens*, is sufficient to reduce PLIN1 binding (Fig. 2i-l and Extended Data Fig. 13). Taken together, our results suggested that sub-optimal phospholipid packing may cause increased turnover of LD-targeting AH proteins and explained the downregulation of AH proteins on LDs from *clstn3b^-/-^* adipocytes.

## CLSTN3B promotes ER/LD contact formation

To understand the mechanistic basis underlying CLSTN3B’s effect of enhancing LD surface structures, we investigated its subcellular localization in adipocytes. We found endogenous CLSTN3B tends to colocalize with the ER marker KDEL around LDs in brown adipocytes (Fig. 3a), likely representing ER/LD contact sites. More prominent CLSTN3B signal was observed around the small LDs emerging after NE treatment (Fig. 3a-b). The LD-associated signals were absent in *clstn3b^-/-^* cells but became stronger in *clstn3b tg* cells (Fig. 3a). Similar observations were made in white adipocytes (Fig. 3a). Immunogold labeling revealed localization of endogenous CLSTN3B to the extensive ER/LD contact in brown adipocytes pre-treated with NE (Fig. 3c), whereas the extent of ER/LD contact is significantly diminished in *clstn3b^-/-^*cells (Extended Data Fig. 14). Electron microscopy also revealed scanty ER/LD contacts on large LDs (Extended Data Fig. 15), consistent with the preferred localization of CLSTN3B to small LDs (Fig. 3b). To better understand CLSTN3B’s preference for small LDs, we performed a NE treatment time-course (Fig. 3d). Before NE-treatment, brown adipocytes contained mostly large LDs with scanty CLSTN3B signal around them. After overnight NE treatment, numerous small LDs emerged with prominent CLSTN3B signal around them, which can be strongly inhibited by the long-chain acyl CoA synthetase inhibitor triacsin C, indicating those small LDs were generated *de novo* from re-esterified free fatty acids released after NE stimulation, consistent with a previous observation^29^. On day 5 after NE withdrawal, all small LDs became large, accompanied by the disappearance of CLSTN3B signal (Fig. 3d). Taken together, we showed that CLSTN3B-mediated ER/LD contact preferentially occurs on newly formed small LDs but becomes profoundly diminished on mature large LDs, explaining why significant downregulation of ER proteins was selectively observed in brown adipocyte relative to white adipocyte LD proteomics (Fig. 2b-c), since small LDs make a substantial contribution in brown adipocytes whereas white adipocytes invariably contain large mature LDs.

**Figure 3.**
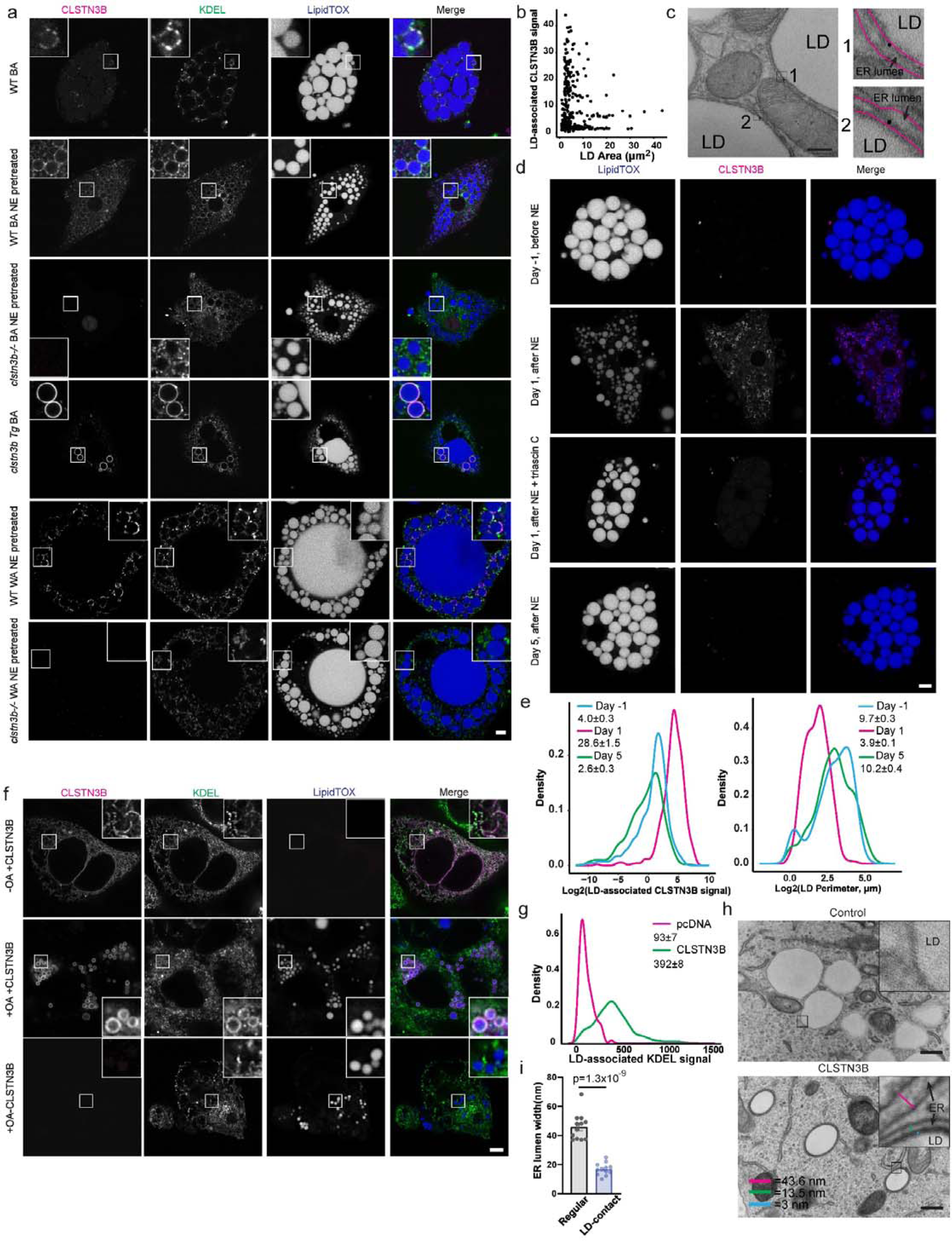
CLSTN3B promotes ER/LD contact formation. **a**, Fluorescence microscopic analysis of CLSTN3B and KDEL localization in primary adipocytes. Scale bar: 5 μm. **b**, Relationship between LD-associated CLSTN3B intensity and LD size. **c**, Electron microscopic analysis of CLSTN3B localization in primary brown adipocytes. Arrows denote gold particles labeling CLSTN3B. Scale bar: 200 nm. **d**, Fluorescence microscopic analysis of CLSTN3B localization in primary brown adipocytes over a NE-treatment time course. Scale bar: 5 μm. **e**, Quantitation of LD-associated CLSTN3B signal intensity and LD size in **d**. **f**, Fluorescence microscopic analysis of CLSTN3B and KDEL localization in HEK293 cells. Scale bar: 5 μm. **g**, Quantitation of LD-associated KDEL signal intensity in HEK293 cells. **h**, Electron microscopic analysis of ER/LD morphology in HEK293 cells -/+ CLSTN3B expression. Scale bar: 400 nm. **i**, Quantitation of the lumen width of regular and LD-associated ER (n=12 measurements).

In heterologous systems, CLSTN3B also induced ER/LD contact as shown by fluorescence and electron microscopy (Fig. 3f-i and Extended Data Fig. 16). Such contacts are extensive (completely encircling LDs), tight (<5 nm distance between the ER and LD membranes), excluding known LD-targeting proteins (PLIN1, PLIN3, CIDEA, and VPS13C), and the width of the ER lumen is narrowed from 46 nm to 17 nm (Fig. 3h-i and Extended Data Fig. 17). Furthermore, CLSTN3B promoted microsome/LD contact in a semi-reconstituted system (Extended Data Fig. 18). We therefore conclude that CLSTN3B is sufficient to promote ER/LD contact formation in non-adipocytes and a cell-free condition.

## CLSTN3B directly bridges ER and LD

To understand the sequence determinants underlying the localization of CLSTN3B, we analyzed its hydrophobicity and predicted its structure with AlphaFold^30^. Such analyses revealed an extended N-terminal hydrophobic domain (approximately 1-170) consisting of multiple helices, a typical transmembrane domain (244-269, identical with the transmembrane domain of CLSTN3), an intervening arginine-rich segment (171-243), and a C-terminal tail (270-357) (Fig. 4a and Extended Data Fig. 19). When expressed in HEK293 cells, the N-terminal fragment (1-198) localizes to the LD but not the ER (Fig. 4b and Extended Data Fig. 20), whereas the C-terminal fragment (199-357) displays a complementary localization pattern. In contrast to full-length CLSTN3B, the LD-localizing N-terminal fragment does not exclude CIDEA (Fig. 4c-d), suggesting that CIDEA exclusion may be caused by the exceptionally tight membrane contact. These data suggest that the N-terminal hydrophobic domain of CLSTN3B targets the LD whereas the C-terminal transmembrane domain is anchored at the ER.

**Figure 4.**
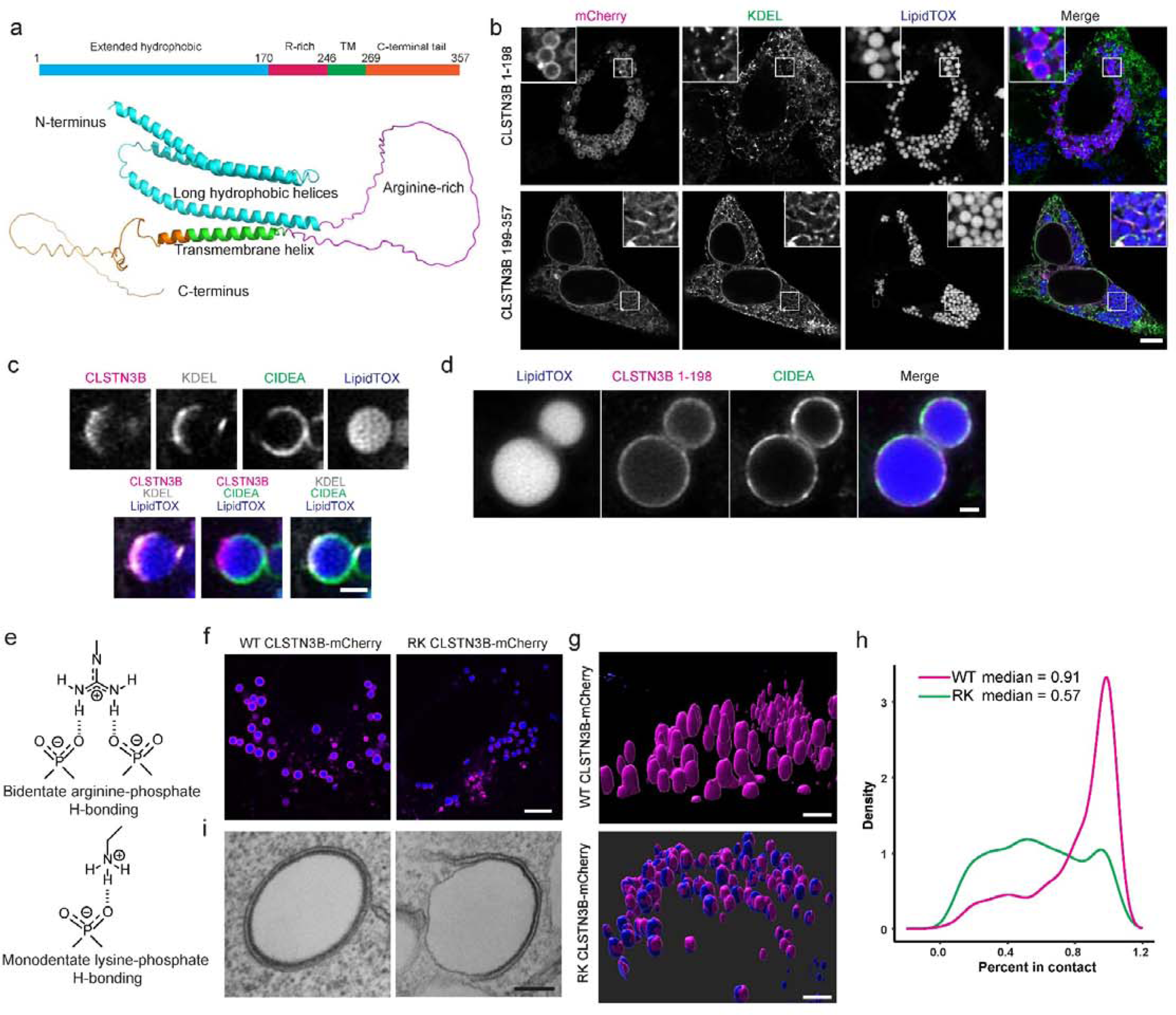
CLSTN3B bridges the LD and the ER. **a**, Ribbon diagram of AlphaFold predicted structure of CLSTN3B. **b**, Fluorescence microscopic analysis of CLSTN3B 1-198 and 199-357 localization in HEK293 cells. **c-d**, Fluorescence microscopic images showing exclusion or non-exclusion of CIDEA by full-length CLSTN3B (**c**) or CLSTN3B 1-198 (**d**). **e**, Illustration of arginine/lysine phosphate hydrogen bonding. **f-i**, Fluorescence microscopic images (**f**), 3D reconstruction (**g**), quantitative analysis of ER/LD contact extent (**h**), and electron microscopic images (**i**) of WT CLSTN3B and RK mutant-mediated ER/LD contact in HEK293 cells. Scale bar: 5 μm in **b** and **f**; 1 μm in **c** and **d**; 3 μm in **g**, and 200 nm in **i**.

The topology of CLSTN3B puts the arginine-rich segment in the space between the LD and the ER. Arginine-rich peptides interact strongly with membranes through bi-dentate hydrogen bonding with phospholipids^31–35^ (Fig. 4e). We therefore hypothesize that the arginine-rich segment could be responsible for the extensive ER/LD contact formation. Indeed, replacing 10 arginine residues with lysine (CLSTN3B RK) significantly reduced the extent of CLSTN3B-induced ER/LD contact formation (Fig. 4f-i). These results thus support a critical role of the arginine-rich segment in inducing extensive ER/LD contact formation.

## CLSTN3B facilitates ER-to-LD phospholipid transfer

Since the ER is the main source of phospholipids for other organelles^36,37^, we hypothesized that CLSTN3B may be promoting ER-to-LD phospholipid flow, explaining why CLSTN3B ablation leads to LD surface structure deficiencies. To test this idea, we first investigated the effect of CLSTN3B on LD surface phospholipid abundance in non-adipocyte cells. LDs from HEK293 cells expressing CLSTN3B displayed 54% higher density of PC and 33% higher density of PE than non-expressing cells, whereas the RK mutant displayed a strongly diminished effect (Fig. 5a).

**Figure 5.**
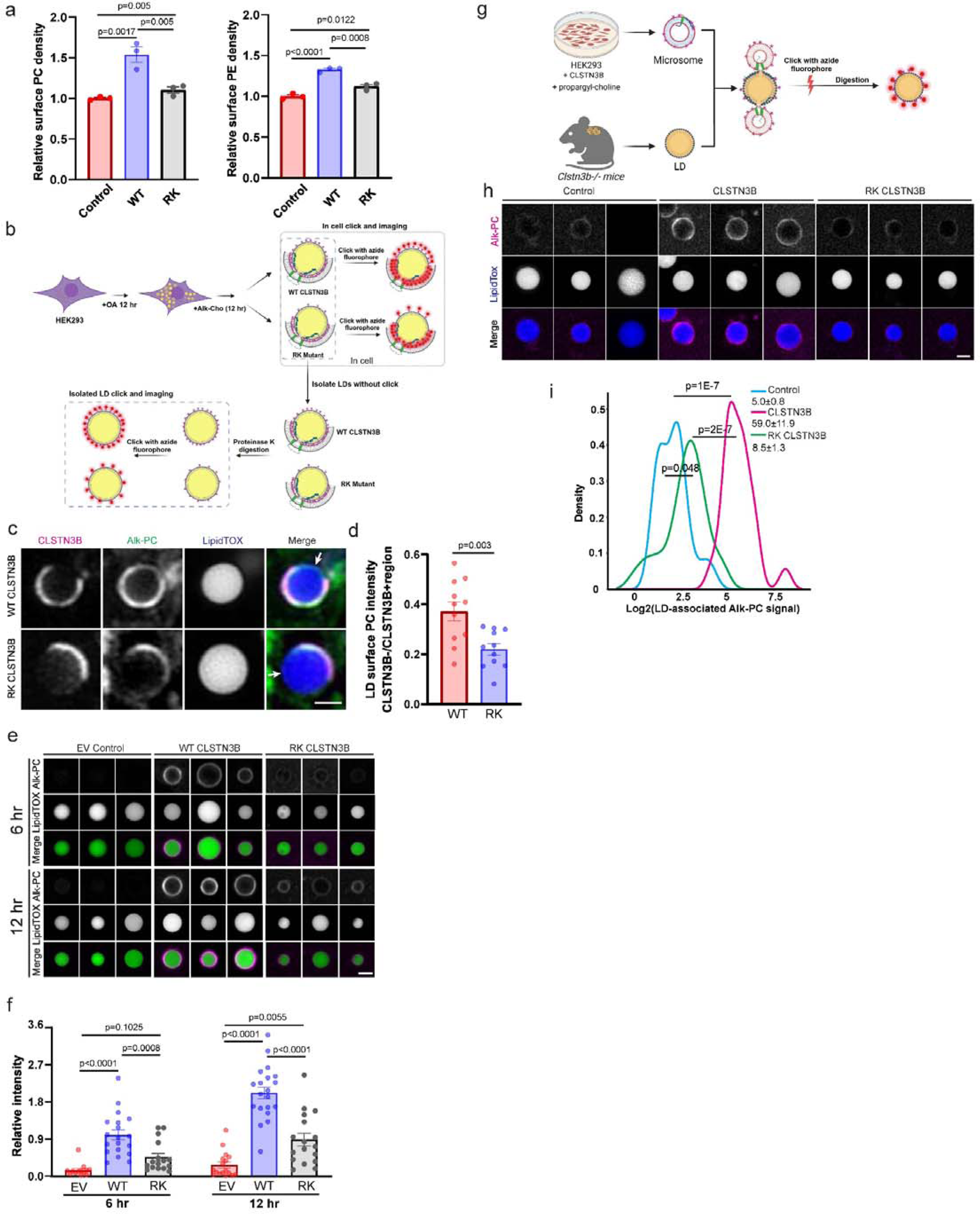
CLSTN3B facilitates ER-to-LD phospholipid transfer. **a**, Fluorometric phospholipid quantitation of LDs isolated from empty vector, WT CLSTN3B or RK mutant-expressing HEK293 cells (n=3 replicates). **b**-**f**, Schematics (**b**), representative images (**c**) and (**e**), and quantitation (**d**) and (**f**) of fixed cell (**c-d**) or isolated LD (**e-f**) click-chemistry based ER-to-LD phospholipid transfer assay. Scale bar: 1 μm. **g-i**, Schematics (**g**), representative images (**h**), and quantitation (**i**) of semi-reconstituted phospholipid transfer assay. Scale bar: 2 μm.

To obtain direct evidence of CLSTN3B-mediated ER-to-LD phospholipid flow, we first induced LD formation in HEK293 cells, then labeled newly synthesized PC with alkyne choline, and finally used an azide-fluorophore to visualize labeled PC (Fig. 5b). We compared the fluorescence level on the part of LD surface free from CLSTN3B versus the part in association with CLSTN3B, as this ratio reflects the extent by which newly synthesized PC have translocated from the ER to LD. The results showed in cells expressing WT CLSTN3B, the ratio between labeled PC on the CLSTN3B-free LD surface to the part associated with CLSTN3B is significantly higher than in cells expressing the RK mutant (Fig. 5c-d), suggesting a larger extent of ER-to-LD phospholipid flow mediated by WT CLSTN3B. To avoid interference from ER phospholipids and exclusively focus on LD monolayer phospholipids, we took an alternative approach of digesting isolated LDs with proteinase K to remove bound ER followed by click-chemistry labeling of newly synthesized PC having translocated from the ER to LD (Fig. 5b and Extended Data Fig. 21). This approach allowed more accurate quantitation of a larger number of LDs and the results again showed that LDs from CLSTN3B-expressing cells displayed significantly higher levels of labeled PC than control cells or cells expressing the RK mutant (Fig. 5e-f), supporting more efficient ER-to-LD phospholipid transfer under CLSTN3B expression. Similar effects could be recapitulated in a semi-reconstituted system: CLSTN3B-containing microsomes promoted much more efficient microsome-to-LD phospholipid transfer than the RK mutant or an empty vector control (Fig. 5g-I and Extended Data Fig. 22). Our data thus support that CLSTN3B facilitates ER-to-LD phospholipids flow, and this process requires optimal ER-LD contact formation driven by the arginine-rich segment.

## The arginine-rich region of CLSTN3B induces membrane hemifusion

We next sought to understand the mechanism of CLSTN3B-mediated ER-to-LD phospholipid flow. This function is unlikely fulfilled by proteins other than CLSTN3B given that the tight ER/LD contact excludes all LD-surface proteins that we have tested. Because arginine-rich peptides are known to promote membrane fusion and phospholipid mixing^32–35^, we hypothesize that CLSTN3B may employ the arginine-rich segment located between the ER and LD membrane to induce fusion between the ER cytosolic leaflet and the LD monolayer and facilitate ER-to-LD phospholipid diffusion. To test this idea, we examined the ability of an CLSTN3B-derived arginine-rich peptide (CLSTN3B 222-232: RTRNLRPTRRR) to promote liposome hemifusion with a FRET-based assay system (Fig. 6a)^38^. We observed that the arginine-rich peptide but not an RK mutant peptide promoted phospholipid mixing (Fig. 6b). The activity of the CLSTN3B-derived arginine-rich peptide exhibited a strong dose dependence on phosphatidic acid (PA) (Fig. 6c), an anionic conical phospholipid favoring negative membrane curvature required for hemifusion stalk formation^39^, but was antagonized by an inverse conical phospholipid lauroyl-lysophosphatidylcholine (LPC) (Fig. 6d), which favors positive membrane curvature and inhibits hemifusion^40^. These results strongly support a direct role of CLSTN3B in promoting ER-to-LD phospholipid diffusion via arginine-rich region-mediated formation of a hemifusion-like structure.

**Figure 6.**
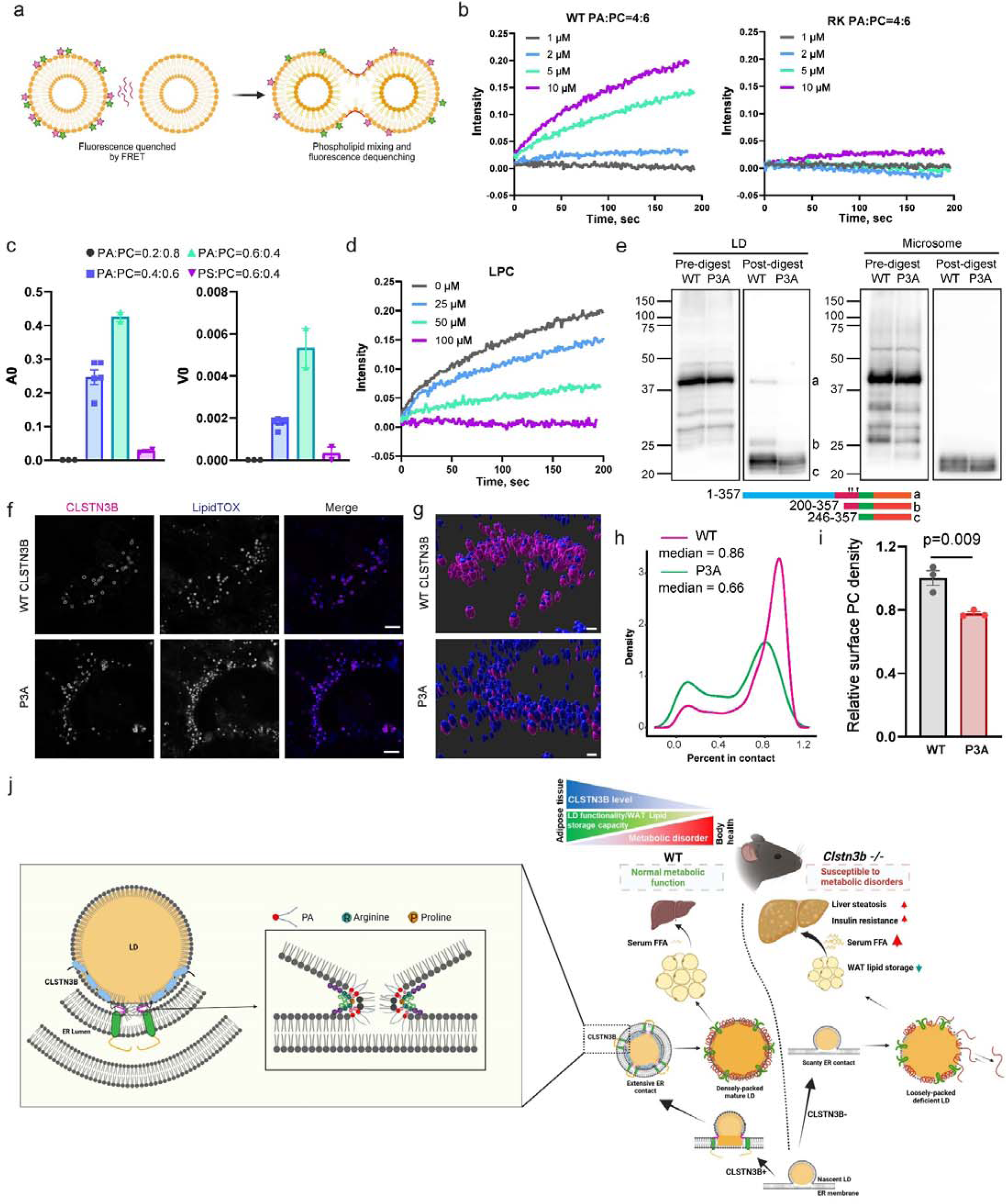
CLSTN3B promotes hemifusion-like structure formation. **a**-**d**, Schematics (**a**) of FRET-based detection of liposome lipid mixing induced by a CLSTN3B-derived R-rich peptide or a RK mutant peptide (**b**), the dependence on phospholipid composition for lipid mixing (**c**), and the effect of LPC on lipid mixing (**d**). **e**, Western blot analysis of WT or P3A CLSTN3B fragments generated from trypsin digestion of LDs from HEK293 cells. The positions of fragments with defined lengths are shown as references (see also Extended Data Fig. 23). Arrows denote positions of mutated prolines. **f-i**, Fluorescence microscopic images (**f**), 3D reconstruction (**g**), quantitative analysis of ER/LD contact extent (**h**), and measurement of LD surface PC levels (**i**) in WT CLSTN3B and P3A mutant-expressing HEK293 cells. Scale bar: 2 μm. **j**, Model of CLSTN3B action at the molecular, organellar, cellular and organismal levels.

Our model for CLSTN3B-supported membrane fusion at the ER/LD contact site predicts that the peptide backbone of the arginine-rich segment needs to sustain abrupt turns in accordance with the negative membrane curvature, given the extremely narrow space at the contact site (Fig. 2h). Interestingly, we noted that proline is the second most abundant amino acid in this region (13%) following arginine (29%) (Extended Data Fig. 23), significantly above its frequency in the mouse proteome (6% and p=0.029 by Chi-square test). To test the significance of the proline residues, we mutated 3 prolines to analines (P220, 228, 241A, or P3A). The P3A mutant was more susceptible to trypsin digestion than the WT CLSTN3B on isolated LDs but not microsomes (Fig. 6e), consistent with the arginine-rich region of the mutant not able to remain tightly associated with the curved membrane surface and hence more accessible for trypsin digestion. Functionally, the P3A mutant is less effective at promoting ER/LD contact formation and enhance LD surface phospholipid levels (Fig 6f-i). Taken together, we have provided evidence supporting CLSTN3B-supported membrane fusion at ER/LD contact, with arginine residues required for interacting with phospholipids and proline residues contributing to the peptide backbone curvature (Fig. 6j).

## Discussion

We have elucidated that the mammalian adipocyte-specific protein CLSTN3B promotes extensive ER/LD contact formation and facilitates ER-to-LD phospholipid diffusion via membrane fusion during the early phase of LD growth, which in turn allows better retaining of LD-targeting proteins as they are recruited to maturing LDs. LDs in adipocytes grow to uniquely large sizes and require a larger supply of surface structural molecules than in other cell types. Hence, CLSTN3B provides an adipocyte-specific mechanism to allow LDs to reach a high level of structural and functional maturity (Fig. 6j). In brown adipocytes, CLSTN3B enhances LD surface structure to reduce surface tension, prevents excessive LD coalescence to ensure sufficient LD surface area, and sensitizes lipolytic response to β-adrenergic stimulation, all beneficial to thermogenesis. In white adipocytes specializing in energy storage, CLSTN3B enhances the surface structure of the large monolocular LDs to protect the internal TG storage from unregulated lipolysis and allow them to undergo expansion in response to caloric overload. Overall, CLSTN3B allows LD size and function to become more polarized tailoring to the needs of distinct cellular contexts. The ability of CLSTN3B to simultaneously promote energy storage and efficient mobilization in response to body needs may have conferred mammals a survival advantage by increasing their adaptability to varying environmental conditions.

Consistent with our finding, a recent study also noted that CLSTN3B deficiency results in enlarged LDs in brown adipocytes, although it was interpreted as excessive *activation* of CIDEA^41^. Our data suggest that insufficient phospholipid supply is likely the main driving force underlying LD coalescence and enlargement, whereas LD-associated CIDEA protein level and activity is *suppressed* in the absence of CLSTN3B, explaining the observation that the largest LDs in *clstn3b^-/-^* brown adipocytes failed to reach comparable sizes as in WT or *clstn3b tg* cells. Importantly, we provide evidence showing that CIDEA, along with multiple known LD surface proteins, is excluded from CLSTN3B-mediated ER/LD contact, strongly suggesting that the function of CLSTN3B at ER/LD contact may not be related to the previously reported CLSTN3B/CIDEA interaction^41^. Nevertheless, our preliminary evidence suggests that the N-terminal LD-targeting domain of CLSTN3B may interact with CIDEA and stabilize its LD-association after LD maturation and ER/LD separation. Further study is required to fully clarify the nature and implication of CLSTN3B/CIDEA interaction.

The selective presence of CLSTN3B and ER contact on small LDs in their early growth phase raises three important questions: how CLSTN3B-mediated ER/LD contact and membrane bridge is initiated with CLSTN3B originally residing in the ER; how the membrane bridge gets resolved to allow ER detachment from mature LDs; whether CLSTN3B continues to facilitate phospholipid replenishment on mature LDs. A recent study showed the membrane fusion machinery induces membrane bridge formation between the ER exit site and LD^15^. We speculate that such membrane bridges could provide an initial route for the N-terminal LD-targeting domain of CLSTN3B to relocate from the ER onto LD surface. Once in place, CLSTN3B can use its unique arginine-rich region to induce additional membrane fusion and promote extensive ER/LD contact. The presence of short forms of CLSTN3B in the ER suggests an intriguing possibility that proteolysis may be involved in the ER/LD separation process as LDs mature, whereas the competition between LD-targeting proteins, such as PLIN1 and CIDEA, and CLSTN3B may also reduce the ER/LD contact area. Lastly, our proteomics results suggest that CLSTN3B may continue to supply phospholipids to mature large LD via ER-derived vesicle or caveolae. Follow-up investigations are required to clarify the role of RABs and corresponding GEFs in supporting CLSTN3B function on large mature LDs.

## Acknowledgement

We thank Dr. Leonid Chernomordik for discussion on the liposome fusion and phospholipid mixing assay; Andrew Lemoff and the Proteomics core at UTSW for the proteomics analysis; the Molecular Pathology core at UTSW for the histology analysis; the EM core at UTSW (supported by NIH grant 1S10OD021685-01A1), especially Dr. Kate Luby-Phelps, Phoebe Doss, and Rebecca Jackson, for project discussion and EM sample preparation; Drs. Tobias Walther and Robert Farese for providing the SEIPIN KO cell line; Drs. Xiaowei Chen and Xinxuan Xu for advice on Click Chemistry experiments; Drs. Xiaowei Chen, Xiaolei Su, Bo Hu, and Rui Chang for critical reading of the manuscript; Xing Zeng is a Rita C. and William P. Clements, Jr. Scholar in Biomedical Research. This study was supported by the Endowed Scholars in Medical Science Program at UTSW, Cancer Prevention and Research Institute of Texas grant RR200084, NIH R01DK135556, and American Heart Association Award 23CDA1050474 to X.Z. The work of K.M. was supported by the Intramural Research Program of the Eunice Kennedy Shriver National Institute of Child Health and Human Development, National Institutes of Health.

## Materials and Methods

### Mouse strains

The *clstn3b^-/-^* and the adipose-specific *clstn3b* transgenic mice were described in a previous publication (Zeng et al, PMID: 31043739). All mice were maintained under a 12 hr light/12 hr dark cycle at constant temperature (23°C or 30°C as specified) with free access to food and water. All animal studies were approved by and in full compliance with the ethical regulation of the Institutional Animal Care and Use Committee of University of Texas Southwestern Medical Center. Male mice of 8-12 weeks of age were used for physiological experiments unless stated otherwise. Sample size was chosen based on literature and pilot experiment results to ensure that statistical significance could be reached. Randomization was not performed because mice were grouped based on genotype.

### Reagents

FFA fluorometric assay kit (Cayman, item no.700310); Lipofectamine 3000 (Invitrogen, L3000-015); Norepinephrine (Sigma, A9512); HCS LipidTOX™ Deep Red Neutral Lipid Stain (Invitrogen™, H34477); HCS LipidTOX™ Green Neutral Lipid Stain (Invitrogen™, H34477); Trypsin Platinum (Promega, VA9000); Protease K (Sigma, P2308); Oleic acid (Sigma, O1383); BSA (Sigma, A7030); FFA-free BSA (Millipore, Code82-002-4); Trypsin inhibitor (Worthington, LS003571); Protease inhibitor cocktail (100X) (Thermo scientific, 1861279); Free glycerol reagent (Sigma, F6428-40ML); Propargyl choline (Cayman, 25870); BTTAA (Click chemistry tools, 1236-100); CalFluor 647 Azide(Click chemistry tools, 1272-1); Phosphatidylcholine Assay Kit (Colorimetric/Fluorometric) (Abcam, ab83377); Phosphatidylethanolamine Assay Kit (Fluorometric) (Sigma, MAK361).

#### Antibody

CLSTN3B antibody was described previously (Zeng et al, PMID: 31043739); KDEL (Abcam, ab184819); PLIN1 (Abcam, ab3526); CGI-58 (Abcam, ab183739); UCP1(Abcam, ab209483); PDI (Cell signaling, C81H6); CS (Abcam, ab129095). Seipin (Cell signaling, 23846S), HA-tag (Cell signaling, 3724S), FLAG (Sigma, F1804), PLIN1 (Cell signaling, 9349S), p-HSL(s660) (Cell signaling, 4126), HSL (Cell signaling, 4107S), Alexa Fluor 546 Fluor Nanogold IgG goat anti-mouse (Nanoprobes, 7410-1ML), Alexa Fluor 488 Fluor goat anti-Rabbit (Invitrogen, A11034), Alexa Fluor 564 Fluor goat anti-mouse (Invitrogen, A11030), Alexa Fluor 405 Fluor goat anti-rabbit (Invitrogen, A31556).

#### Constructs

PCDNA3.1-*clstn3b* was described previously^15^; *clstn3b*-mCherry, *clstn3b*(1-131)-mCherry, *clstn3b*(1-198)-mCherry, *clstn3b*(131-357)-mCherry, *clstn3b*(199-357)-mCherry, *seipin*, *Clostridium perfringensfrom* alpha toxin, and *clstn3b-C1-FLAG* were synthesized by Gene Universal Inc. (Newark DE 19713); BFP ER-reporter was purchased from Addgene (49150).

### Electron microscopy

Primary adipocytes were isolated as previously described and allowed to attach to MatTek glass bottom grid dishes (P35G-1.5-14-CGRD) (Zeng et al, PMID: 31043739). The cells were treated with 1 μM norepinephrine overnight to induce the formation of small LDs when specified. Cells were then fixed with 4% paraformaldehyde + 0.1% glutaraldehyde in PBS containing 7.5% sucrose for 30 min at RT, blocked with 50 mM glycine for 15 min, permeabilized with 0.25% saponin for 30 min. Cells were than incubated with the CLSTN3B antibody at 4°C overnight and then with the fluoronanogold secondary antibody (Nanoprobes 7403-1) for 2 h at RT. Images for taken on a Zeiss 900 confocal microscope to identify the coordinates of the cells for EM analysis. The cells were further gold-enhanced for 2.5 min using a gold enhancement kit (Nanoprobes). After washing with water and 0.1M cacodylate buffer, samples were fixed with 1% OsO4 and 0.8% potassium ferricyanide in 0.1M cacodylate buffer for 1h, stained en bloc with 2% aquaous uranyl acetate, dehydrated with increasing concentration of ethanol, and embedded in Epon. Blocks were sectioned with a diamond knife (Diatome) on a Leica Ultracut 7 ultramicrotome (Leica Microsystems) and collected onto copper grids. Images were acquired on a JOEL 1400 Plus transmission electron microscope equipped with a LaB6 source using a voltage of 120 kV, and the images were captured by an AMT BIOSPRINT 16M-ActiveVu mid mound CCD camera.

### Fixed cell imaging

Primary adipocytes were isolated as previously described and attached to glass coverslips pre-coated with laminin and poly-D-lysine (Zeng et al, PMID: 31043739). HEK293 cells and U2OS cells were plated onto glass coverslips and treated with 400 μM oleic acid for 12-24 hr to induce LD formation. Images were taken on a ZEISS 900 confocal microscope with Airyscan.

### Image analysis

To analyze isolated lipid droplets, images were imported to Imaris (Bitplane). Individual LDs were rendered by the “Surface” function and surface area and volume were measured. The rendering parameters were kept the same between treatments in each experiment. To analyze the extent of CLSTN3B wrapping of LDs, individual LDs and the surrounding CLSTN3B signals were rendered by the “Surface” function. The fraction of the volume bound by the LD surface overlapping with the volume bound by the CLSTN3B signal surface was calculated with the “Object-Object statistics” function. To analyze LD-associated signals (PC, microsomes, KDEL, CLSTN3B or PLIN1), images were imported to ImageJ (NIH). Individual LDs were detected by the “Analyze particle” function, and a larger ROI was generated for each LD via the “Dilate” function to encompass surrounding signal of interest. Fluorescence signal intensity associated with LDs was then measured, normalized or transformed, and plotted. To analyze ER wrapping of LD in EM images, the LD perimeter and the fraction in contact with the ER are selected and calculated with the polygon tool in Image J. To analyze ER lumen width in EM images, a line is drawn through the lumen and perpendicular to the ER membrane and the length of the line is calculated in Image J. All plots were generated in R.

### LD isolation and digestion

For BAT LD, BAT was dissected, minced into tiny pieces with a spring scissor and transferred to a motorized homogenizer in HES buffer (20mM HEPES + 1mM EDTA + 250mM Sucrose). The homogenate was filtered with double-layer gauze and centrifuged at 2000 g for 5 min. The infranatant was removed with a syringe and the buoyant LD fraction was transferred with a wide-opening tip into 5 ml tubes and washed with HES buffer 2 times. The LD was then transferred to Ultra-Clear ultracentrifuge tubes (Beckman-Coulter), adjusted to a final concentration of 20% sucrose, and overlaid by 5% sucrose/HE and HE (20mM HEPES + 1mM EDTA). The gradient was centrifuged at 16,000 g for 10 min at 4℃. The buoyant LD was then collected.

For WAT LD, WAT was dissected, minced into tiny pieces with a spring scissor and transferred to digestion buffer. Digest on a rotator at 37LJ for 30min. Filter with 100um cell strainer into a 50mL tube. (Digest buffer: HBSS + 2mg/ml Collagenase B + 1mg/ml Trypsin inhibitor + 4% BSA (FA free) + 25U/ml Benzonase (1:10000)). Transfer the cell suspension to a 5 mL tube and wash with HES buffer 3 times. Disrupt the cell in HES buffer by gently passing through a 25G needle with 10 times. Wash 3 times with HES buffer. Don’t centrifuge or vortex, let it stand for 10 minutes until the buoyant LD fraction float to the upper layer. Keep the buoyant LD fraction. For measuring the PL of WAT LD, transfer the LD fraction to Ultra-Clear ultracentrifuge tubes (Beckman-Coulter), diluted to a final concentration of 20% percoll. Centrifuge 10000g X 20min at 4LJ, keep the buoyant LD fraction.

For HEK293 LD, 15-cm dishes of HEK293 cells were grown to 90% of confluence and treated with 400 μM oleic acid in growth medium for 24 h. Cells were scraped and collected in 2 mL of lysis buffer (25 mM Tris–HCl, pH 7.4, 100 mM KCl, 1 mM EDTA, 5 mM EGTA, and protease inhibitor cocktail), and lysed by passing through a needle (27-gauge). The lysates were then centrifuged at 1,500 g for 5 min. The supernatants were adjusted to 2.5 mL final volume containing 20% sucrose and transferred into a 10 mL polycarbonate ultracentrifuge tube, and overlaid sequentially with 2.5 mL of 10% sucrose, 2.5 mL of 4.2% sucrose, and 2.5 ml of lysis buffer. The gradient was centrifuged at 150,000 g for 1 hr at 4℃. The buoyant LD was then collected.

For topology determination, LDs isolated from BAT were incubated with trypsin (50 μg/ml, Promega) for 30 min, followed by centrifuging LD fractions through the aforementioned sucrose gradient at 210,000g for 1 hr. The buoyant LD fraction and the pellet fraction were collected separately.

To remove the LD-bound organelles, the LD fraction was digested with 1 mg/mL proteinase K for 5-15 min at 37 ℃ and 1mM PMSF was added to inactivate proteinase K. The mixture was centrifuged through the aforementioned sucrose gradient at 210,000g for 1 hr. The buoyant LD was then collected.

### Immunoprecipitation

HEK293 cells were transfected with the indicated plasmids After 48 hours, cells were washed once with PBS, resuspended in lysis buffer (25 mM Tris-HCl pH 8.0, 150 mM NaCl and 1× protease inhibitor cocktail), and homogenized with a motorized homogenizer. The lysate was then supplemented with 1% digitonin, incubated on rotator at 4°C for 2 hr, and centrifuged at 20,000g for 15 min at 4 °C. FLAG-tagged protein was isolated with anti-FLAG M2 magnetic beads (Sigma, M8823) for 2 hr at 4 °C and eluted with 100 μg/ml 3x FLAG peptide. **Proteomics.** LD was isolated from mouse BAT or WAT as described above without protease digestion. The suspension was mixed with 10x volume of acetone and incubated at −20℃ overnight for delipidation and protein precipitation. The mixture was centrifuged at 12,000 g for 5 min. The pellets were washed with acetone and dried by heating at 60℃ for at least 15 min. Pellets were then washed with 20% TCA to remove excess sucrose, washed with acetone, and dried. The pellets were dissolved in 50 mM triethylammonium bicarbonate (TEAB, pH=8), 5% SDS at 60℃ with shaking for 30 min. Protein concentration was determined with a BCA method. Proteomics analysis was performed following a previously described procedure (Cui et al, PMID: 34489413)

### Phospholipids quantitation

LD was isolated from mouse BAT, WAT or HEK293 cells and digested with proteinase K following the procedure described above. The collected LD fractions (100-200 μL suspension in HES buffer) were transferred into round bottom glass tubes and extracted with a mixture of 1 mL hexane, 1 mL methyl acetate, 0.75 mL acetonitrile and 1 mL water as previously described. The extracts were vortexed for 5 s and centrifuge at 2,671 g for 5 min to partition into 3 phases. The upper and middle phases were collected into separate glass tubes and dried under N_2_. Phospholipids in the middle phase were measured with the Phosphatidylcholine Assay Kit (Colorimetric/Fluorometric) (Abcam, ab83377) and Phosphatidylethanolamine Assay Kit (Fluorometric) (Sigma, MAK361) following the manufacturer’s instructions. TG in the upper phase was dissolved in 350 μL ethanolic KOH (2 part EtOH and 1 part 30% KOH) and incubated overnight at 55°C for complete hydrolysis. The volume was then brought to 1200 μL with H2O: EtOH (1:1) and vortexed to mix. Two hundred μL was transferred to a new tube, mixed with 215 μL 1M MgCl_2_ and vortexed. The mixture was incubated on ice for 10 min and centrifuged at 13,000g for 5 min. Glycerol content in the supernatant was determined with the Free Glycerol Reagent (Sigma, F6428) following the manufacturer’s instructions.

For phospholipidomics analysis, a 50 µL aliquot of LD suspension was transferred to fresh glass tubes for liquid-liquid extraction (LLE). LLE were performed at room temperature (including centrifugation) to maintain consistent solubility and phase separation. For the three-phase lipid extractions (3PLE), 1 mL of hexanes, 1 mL of methyl acetate, 0.75 mL of acetonitrile, and 1 mL of water were added to the glass tube containing the sample. The mixture was vortexed, then centrifuged at 2,671×g for 5 min, resulting in separation of three distinct liquid phases. The middle organic phase (polar lipid content), was collected in a separate glass tube with a Pasteur pippete and spiked with 20 μL of a 1:5 diluted SPLASH Lipidomix standard mixture. The samples were dried under N_2_ air flow and resuspended in 400 µL of hexane. Lipids were analyzed by LC-MS/MS using a SCIEX QTRAP 6500+ (SCIEX, Framingham, MA) equipped with a Shimadzu LC-30AD (Shimadzu, Columbia, MD) high-performance liquid chromatography (HPLC) system and a 150×2.1 mm, 5 µm Supelco Ascentis silica column (Supelco, Bellefonte, PA). Samples were injected at a flow rate of 0.3 mL/min at 2.5% solvent B (methyl tert-butyl ether) and 97.5% Solvent A (hexane). Solvent B was increased to 5% over 3 min and then to 60% over 6 min. Solvent B was decreased to 0% during 30 sec while Solvent C (90:10 (v/v) isopropanol-water) was set at 20% and increased to 40% during the following 11 min. Solvent C is increased to 44% over 6 min and then to 60% over 50 sec. The system was held at 60% solvent C for 1 min prior to re-equilibration at 2.5% of solvent B for 5 min at a 1.2 mL/min flow rate. Solvent D [95:5 (v/v) acetonitrile-water with 10 mM Ammonium acetate] was infused post-column at 0.03 ml/min. Column oven temperature was 25°C. Data was acquired in positive and negative ionization mode using multiple reaction monitoring (MRM). The LC-MS/MS data was analyzed using MultiQuant software (SCIEX). The identified lipid species were normalized to its corresponding internal standard. All solvents used were either HPLC or LC/MS grade and purchased from Sigma-Aldrich (St Louis, MO, USA). Splash Lipidomix stadndards were purchased from Avanti (Alabaster, AL, USA). All lipid extractions were performed in 16×100mm glass tubes with PTFE-lined caps (Fisher Scientific, Pittsburg, PA, USA). Glass Pasteur pipettes and solvent-resistant plasticware pipette tips (Mettler-Toledo, Columbus, OH, USA) were used to minimize leaching of polymers and plasticizers.

To calculate LD surface phospholipids density, we divided phospholipids abundance as measured by the fluorometric kit or phospholipidomics by total LD surface area of each sample. To calculate total LD surface area, we divided total TG content of each sample by the mean LD volume to derive total LD number, which is then multiplied by the mean LD surface area. Mean LD volume and surface area were determined by analysis of LD images as described above.

### *In vitro* reconstitution of CLSTN3B-mediated ER/LD contact formation and phospholipids transfer

To reconstitute CLSTN3B-mediated ER/LD contact formation, HEK293 cells were transfected with the *clstn3b*-mCherry or the BFP-ER reporter and cultured for 48 hr. Then cells were washed with warm PBS twice, scraped off, and lysed by passing through a 27G needle 10 times. The lysates were centrifuged at 1000 g for 10 min to remove nuclei and cell debris. The supernatants were centrifuged through the aforementioned sucrose gradient at 210,000 g for 1 hr. The microsome pellets were suspended in the assay buffer (25 mM HEPES, 150 mM NaCl, 10 mM MgCl_2_, 1 mM CaCl_2_,1 mM ATP, 0.5 mM GTP). LDs were harvested from and a parallel batch of HEK293 cells treated with 400 μM oleic acid in growth medium for 24 hr as described above. LDs and microsome suspension were then mixed and incubated at 37°C overnight. The mixture was stained with HCS LipidTOX™ Green Neutral Lipid Stain (1:200) for 15 min and imaged on a ZEISS 900 confocal microscope. To reconstitute CLSTN3B-mediated phospholipids transfer, HEK293 cells were transfected with the *clstn3b*-mCherry, RK-*clstn3b*-mCherry, SEIPIN-HA or the BFP-ER reporter construct for 24 hr and then treated with 100 μM propargyl-choline for 12-24 hr. Microsome suspension was prepared as described above. LDs isolated from *clstn3b*^-/-^ mouse BAT was incubated with the microsome suspension on a RotoFlex (Argos, R2200) set at a low speed at RT for 12hr or overnight. The mixture was then centrifuged at 100 g for 5min. The buoyant LD fraction was collected and incubated with the click reaction solution (50 μM Calfluo647 azide, BTTAA-CuSO4 complex (50 mM CuSO4, BTTAA/CuSO4 6:1, mol/mol) and 2.5 mM sodium ascorbate) for 1 hr at RT. The LDs were then digested with 1 mg/mL protease K for 5 min, treated with 1 mM PMSF for 5 min, and centrifuged at 100 g for 5 min. The buoyant LD fraction was stained with HCS LipidTOX™ Green Neutral Lipid Stain (1:200) for 15 min. Images were then taken on a ZEISS LSM 900 confocal microscope.

### *In vivo* ER-to-LD phospholipid transfer assay

HEK293 cells were transfected with WT-CLSTN3B-mCherry, RK-CLSTN3B-mCherry, or an empty pCDNA3.1 vector for 24 hrs and then treated with 400 μM oleic acid in growth medium for 24 hrs. To label newly synthesized PC, the growth medium was supplemented with 200 μM propargyl-choline for 6 and 12 hrs, respectively. Lipid droplets were isolated at those time points, digested with proteinase K to remove bound ER, labeled with Calfluo647 azide, and imaged as described above.

### Lipolysis assay

Lipolysis assay on freshly isolated primary white and brown adipocytes were performed as previously described (Chen et al, PMID: 28988768). To measure the whole-body response to CL-316,243, mice were put under isoflurane anesthesia and i.p. injected with CL-316,243 at 1 mg/kg. Blood samples were collected from tails at 0- and 10-min. Serum FFA levels were measured with the Free fatty acid fluorometric assay kit (Cayman, 700310) following the manufacturer’s instructions.

### Liposome preparation

Large unilamellar vesicles (LUV) were prepared as described before (Yang et al., Biophys. J., 2010). Lipid mixtures dissolved in a benzene/methanol mixture (9:1 ratio) were frozen in liquid nitrogen and freeze-dried overnight in a CentriVap vacuum concentrator (Labconco, USA). The dried lipid was then resuspended in a buffer (100 mM NaCl, 10mM Hepes, 5 mM EGTA, pH 7.0) at total lipid concentration of 1 mM. Subsequently, the lipid suspension underwent 10 freeze-thaw cycles by alternating immersion in liquid nitrogen and a 50°C water bath. Finally, the suspension was passed ten times through the double-stacked nucleopore polycarbonate track etch membrane filters with a nominal pore size of 0.1 µm using a Lipex liposome extruder.

### Lipid mixing experiments

Lipid mixing was measured by the release of fluorescence resonance energy transfer between TopFluor PE (0.5 mol %) and rhodamine PE (0.25 mol %). Unlabeled liposomes of various lipid compositions and liposomes labeled with a self-quenching concentration were added to 2 mL of buffer in quartz cuvette in a ratio of 10:1 to a total lipid concentration of 10 μM. Different concentrations of peptides were then added to induce lipid mixing between liposomes. Lipid mixing between labeled and unlabeled liposomes results in a dilution of fluorescent probes and an increase in dye fluorescence due to a relief of self-quenching. Fluorescence changes induced by peptides were recorded under constant stirring using a PC1 photon-counting spectrometer (ISS, USA) with λex = 480 nm and λem = 505 nm. At the end of each recording, complete dequenching of the dye was induced by adding Triton X-100 (0.1% v/v final concentration). All experiments were performed at 37°C. The degree of dye quenching was calculated as Q(t) = 100 × (F(t) - F0)/(Ftriton - F0), where F(t), F0, and Ftriton are fluorescence at time t, before the addition of peptide and after addition of Triton X-100, respectively. The initial rate (v0) and final extent (A0) of lipid mixing were estimated from the fit of the Q(t) curve by the expression Q(t) = A0 - A0* exp(-t/τ) with v0 = A0/τ.

### PLIN1 AH peptide competition assay

PLIN1 AH peptides (human PLIN1 108-143: PPEKIASELKDTISTRLRSARNSISVPIASTSDKVL, FITC-modified or unmodified) were synthesized by ThermoFisher and dissolved in the emulsion buffer (20 mM Tris-HCl, 150 mM NaCl, pH 8.0) at a concentration of 1 mg/mL. Phospholipid (L-a-Phosphatidylcholine soybean, P7443, Sigma) was dissolved in chloroform at a concentration of 32 mM as a stock solution. A 5 μL aliquot of the phospholipid stock solution was dried under nitrogen stream in a 1.5 mL microcentrifuge tube, followed by the addition of 90 μL emulsion buffer, 5 μL Triolein (>99% purify, T7140 Sigma), and 5 μL PLIN1-AH-FITC. For neat oil droplets, 90 μL emulsion buffer was mixed with 5 μL triolein and 5 μL PLIN1-AH-FITC in a clean 1.5 ml microcentrifuge tube. The tubes were then vortexed manually at a fixed angle of ∼30° for 10 cycles of 30 s on/30 s off at 25°C and sonicated in a Bioruptor® Pico sonication device (B01080010) for 30s at a medium frequency level to allow emulsion to happen. To start the competition assay, the emulsified neat oil droplets/phospholipid-coated droplets were mixed with an equal volume of unlabeled PLIN1 AH peptide solution to achieve a 20-fold excess of unlabeled PLIN1 AH peptide over FITC-labeled peptide. Droplet images were then captured at indicated time points with a Zeiss LSM900 confocal system and analyzed with ImageJ.

### PLIN1-mCherry FRAP

Isolated primary brown adipocytes from BAT were cultured for two days and infected with a PLIN1-mCherry adenovirus for 24 hours. Regions of interest were selected to ensure that one entire LD was included to avoid rapid recovery due to intra-LD exchange and then bleached by 100% laser power (546 diode laser), followed by time-lapse scanning with a 5-minute interval for a total 120 minutes. Fluorescence intensity of PLIN1-mcherry on LD surface was quantitated by ImageJ and calculated as the percentage of the pre-bleach value.

### Cycloheximide chase assay

Isolated stromal vascular fraction (SVF) from the BAT of 6-8 week old WT or *clstn3b^-/-^* mice were seeded into 6-well plates and induced to differentiate following a previously published procedure (DMEM + 10% FBS + 1x penicillin/streptomycin and supplemented with 20 nM insulin, 1 mM dexamethasone, 0.5 mM isobutylmethylxanthine, 1 nM rosiglitazone, and 1 nM 3,3,5-triiodo-L thyronine for 2 days). Cells were then kept in a maintenance medium (DMEM + 10% FBS + 1x penicillin/streptomycin supplemented with 20 nM insulin and 1 nM rosiglitazone). On Day 6 post induction, cells were treated 100 μg/mL cycloheximide or vehicle in the maintenance medium for indicated time periods and protein was extracted for immunoblotting.

### LD-targeted alpha toxin assay

*Clostridium perfringens* alpha toxin (Uniprot accession ID: P0C216) was targeted to LD by fusing with the N-terminal LD-targeting domain of CLSTN3B (1-198) and mCherry (CLSTN3B 1-198/mCherry/AT). A catalytically inactive mutant was constructed by introducing two mutations (D84N and H164S). HEK293 cells were co-transfected with the alpha toxin and a PLIN1-mclover3 construct in, cultured for 24 hours and then treated with OA for 12 hours to induce LD formation. LDs were then isolated and washed twice with PBS (Mg^2+^ and Ca^2+^ free). Alpha toxin activation was achieved by adding 1 mM ZnCl_2_ and CaCl_2_ followed by incubation at 37°C for 1h. The mixture was then centrifuged at 17000 g centrifuge for 20 min to isolate LDs. Separate LD aliquots were taken for fluorescence imaging, phospholipid quantitation, and Western blot analysis.

### Real-time qPCR analysis

The following primers were used for qPCR analysis of gene expression. *Cd36*-fwd, GGACATTGAGATTCTTTTCCTCTG, rev, GCAAAGGCATTGGCTGGAAGAAC; *clstn3b*-fwd, CTCCGCAGGAACAGCAGCCC, rev, AGGATAACCATAAGCACCAG; *tnf*-fwd, GGTGCCTATGTCTCAGCCTCTT, rev, GCCATAGAACTGATGAGAGGGAG; *ccl2*-fwd, GCTACAAGAGGATCACCAGCAG, rev, GTCTGGACCCATTCCTTCTTGG; *adgre1*-fwd, CGTGTTGTTGGTGGCACTGTGA, rev, CCACATCAGTGTTCCAGGAGAC; *lipe*-fwd, TTGGGGAGCTCCAGTCGGA, rev, TCGTGCGTAAATCCATGCTGT; *fabp4*-fwd, ACACCGAGATTTCCTTCAAACTG, rev, CCATCTAGGGTTATGATGCTCTTCA; *L19*-fwd, GGTCTGGTTGGATCCCAATG, rev, CCCATCCTTGATCAGCTTCCT; *lpl*-fwd, GGGAGTTTGGCTCCAGAGTTT, rev, TGTGTCTTCAGGGGTCCTTAG; *cfd*-fwd, CATGCTCGGCCCTACATGG, rev, CACAGAGTCGTCATCCGTCAC; *cidec*-fwd, TCGGAAGGTTCGCAAAGGCATC, rev, CTCCACGATTGTGCCATCTTCC; *bscl2*-fwd, GTCTGTGTTCCTCTATGGCTCC, rev, CCAGTGAGACATTGGCAACAGG; *abhd5*-fwd, AGATGTGCCCTCAGGTTGGACA, rev, ATCTGGTCGCTCAGGAAAACCC; *plin1*-fwd, TACCTAGCTGCTTTCTCGGTG, rev, GTGGGCTTCTTTGGTGCTGT; *cidea*-fwd, ATCACAACTGGCCTGGTTACG, rev, TACTACCCGGTGTCCATTTCT.

### Statistical analysis

All data shown are mean ± SEM. Statistical significance was calculated by unpaired Student’s two-sided t-test for comparisons between two groups and one-way ANOVA with Tukey’s post hoc test for comparisons between three groups. Wilcoxon test was used for nonparametric test between samples deviating from normal distribution. Two-way Repeated Measurement ANOVA was used to analyze glucose tolerance test data. All experiments have been successfully repeated with similar results for at least three times.

## Supplementary figure legends are provided in the figures

**Supplementary Table 1.**

Proteomics analysis of brown and white adipocyte LD isolated from WT and *clstn3b^-/-^* mice (n=4 mice).

**Supplementary Table 2.**

Mass spectrometry analysis of CLSTN3B immunoprecipitation from HEK293 cells.

---

**Extended Data Fig 1.**
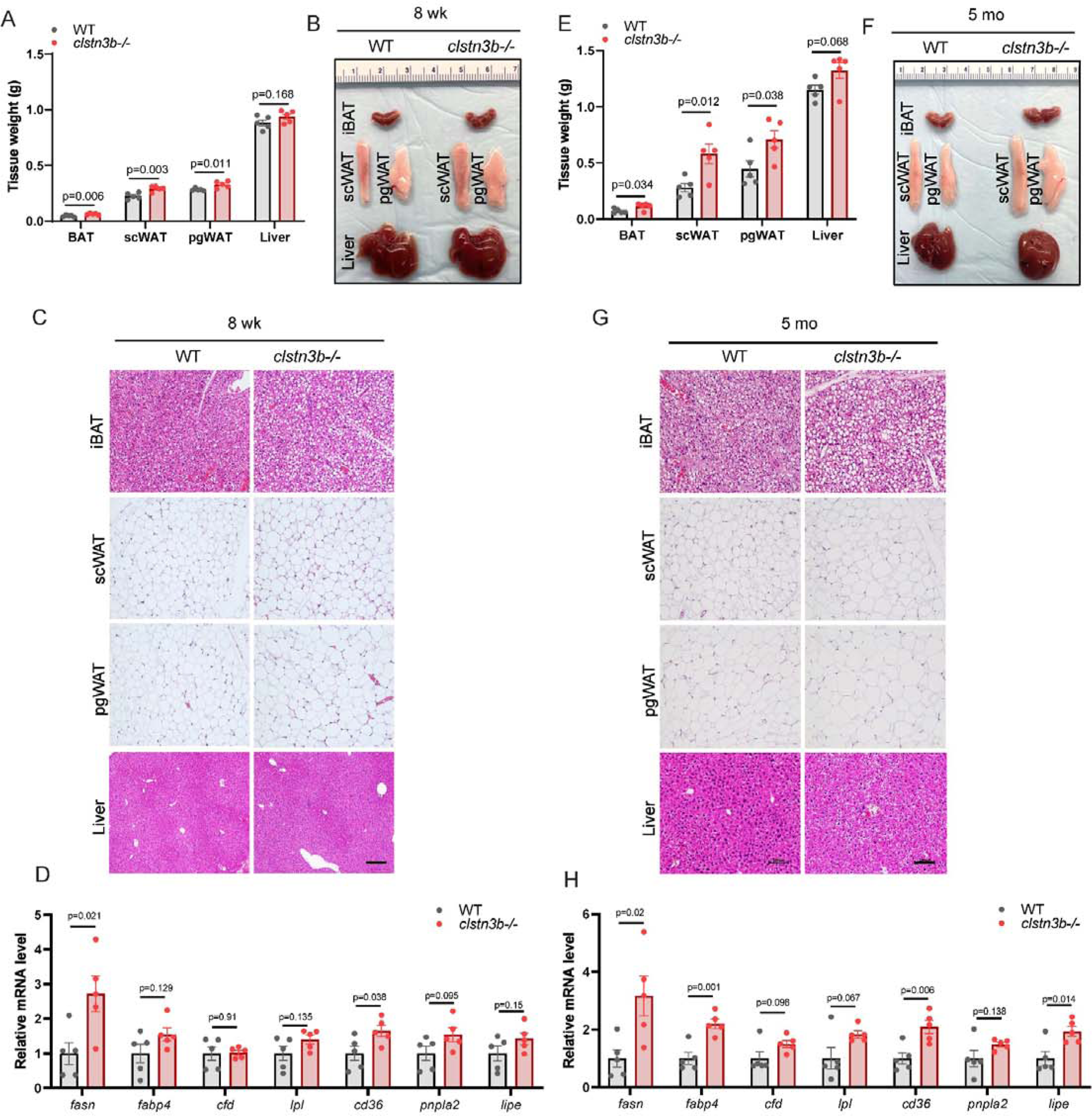
CLSTN3B ablation does not cause defective adipogenesis. A-H, Tissue weights (A, E), gross appearances (B, F), iBAT, scWAT, pgWAT, and liver histology (C, G), and pgWAT gene expression analysis (D, H) of 8 week-old (A-D) or 5 month-old (E-H) WT and *clsfn3b·*^1^*•* mice on chow diet at room temperature (n=6 mice).

**Extended Data Fig 2.**
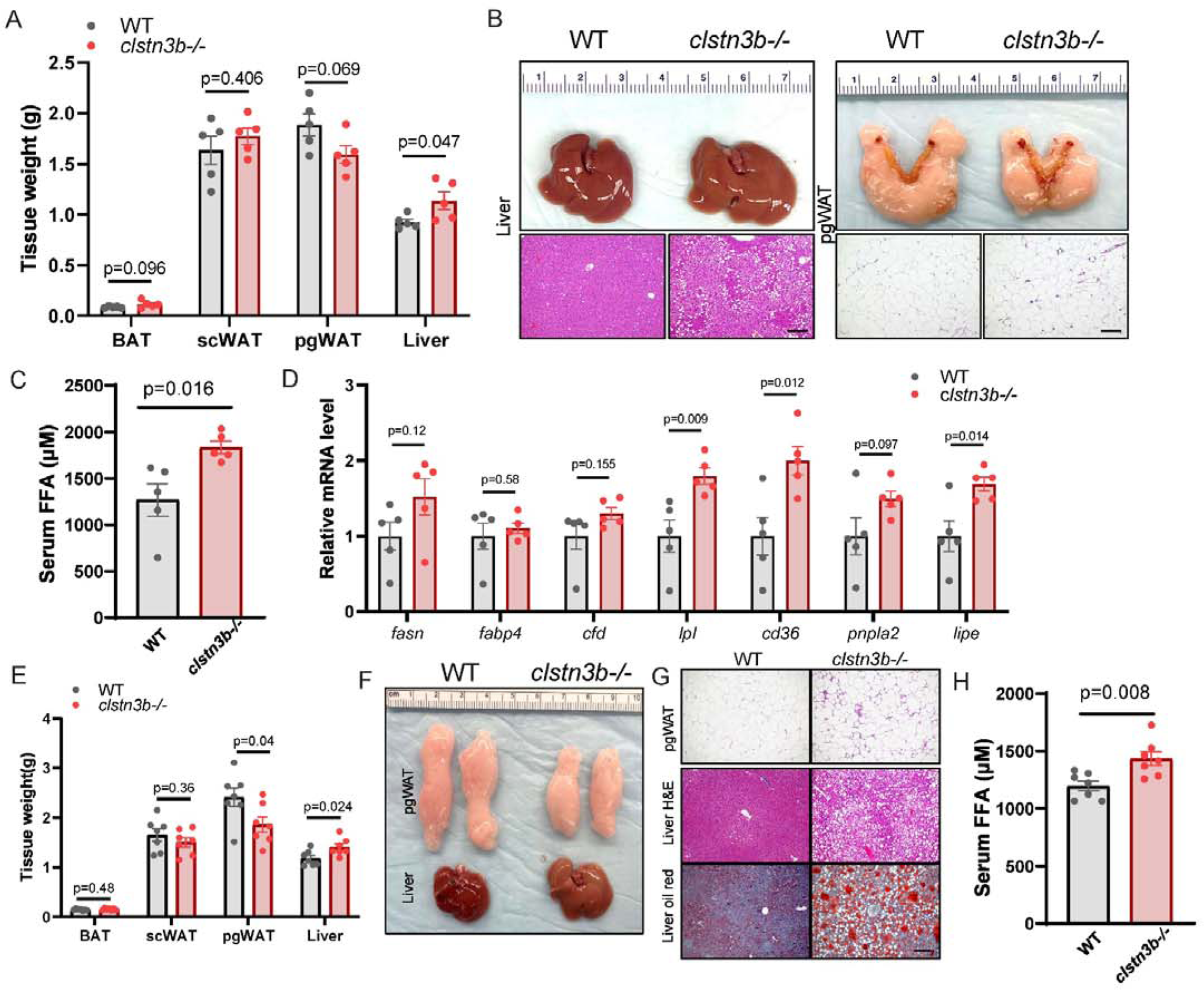
CLSTN3B enhances lipid storage in adipocytes. A-D, Tissue weights (A), gross appearances, pgWAT and liver histology (B), serum FFA (C), and pgWAT gene expression analysis (D) of WT and *clstn3b·*^1^*•* female mice on HFD at thermoneutrality (n=5 mice). E-H, Tissue weights (E), gross appearances (F), pgWAT, liver histology and liver oil red staining (G), serum FFA (H) of WT and *clstn3b-1-* female mice on HFD at room temperature (n=7 mice). Scale bar: 100 µm.

**Extended Data Fig 3.**
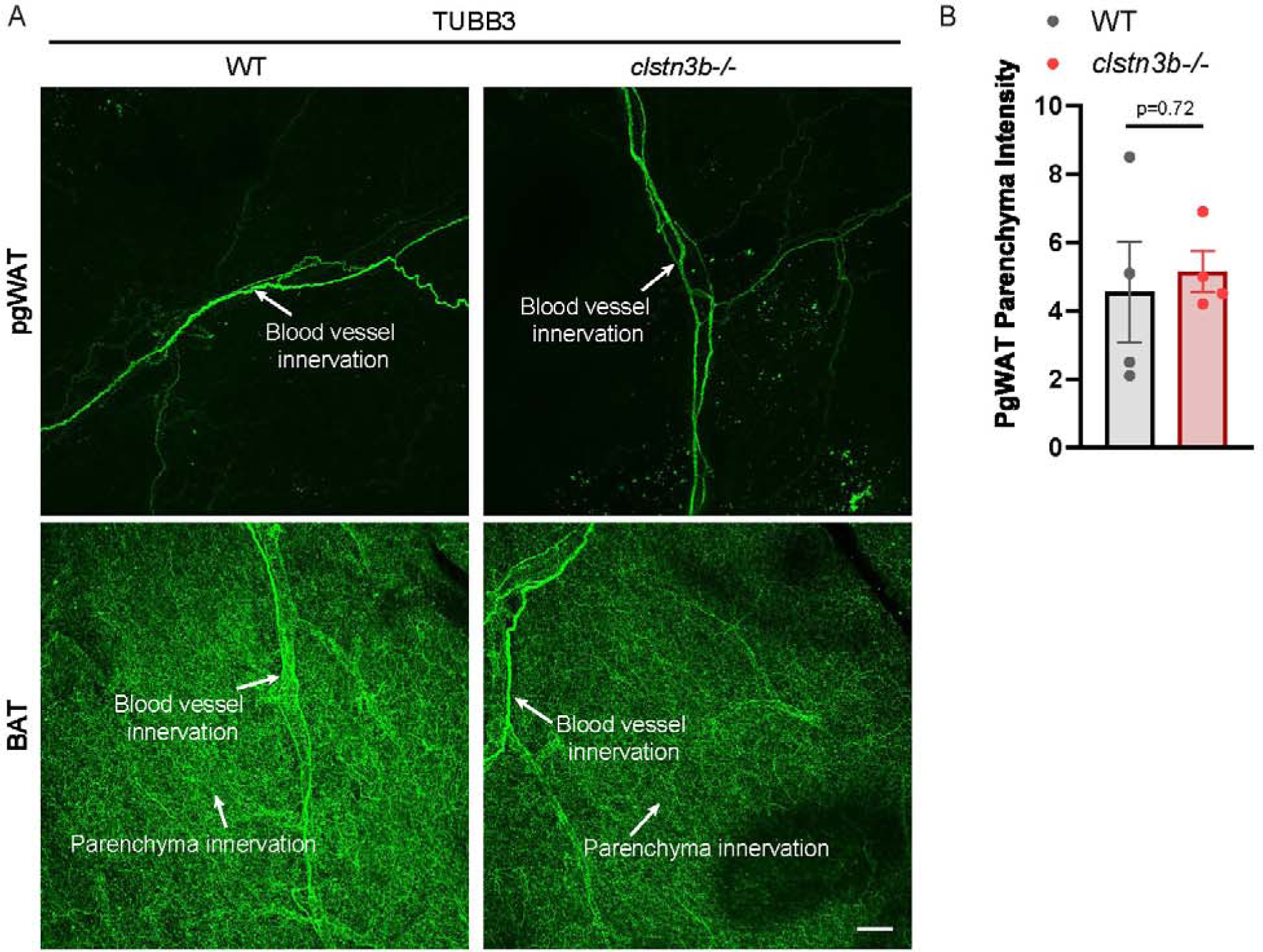
CLSTN3B ablation does not affect sympathetic innervation of the pgWAT. A, TUBB3 immunostaining of pgWAT and BAT from WT and *clstn3b-*^1^*-* mice. Note the different levels of sympathetic innervation in pgWAT and BAT; B, Quantitation of TUBB3 signals in the pgWAT parenchyma. Sea le bar: 100 µm.

**Extended Data Figure 4.**
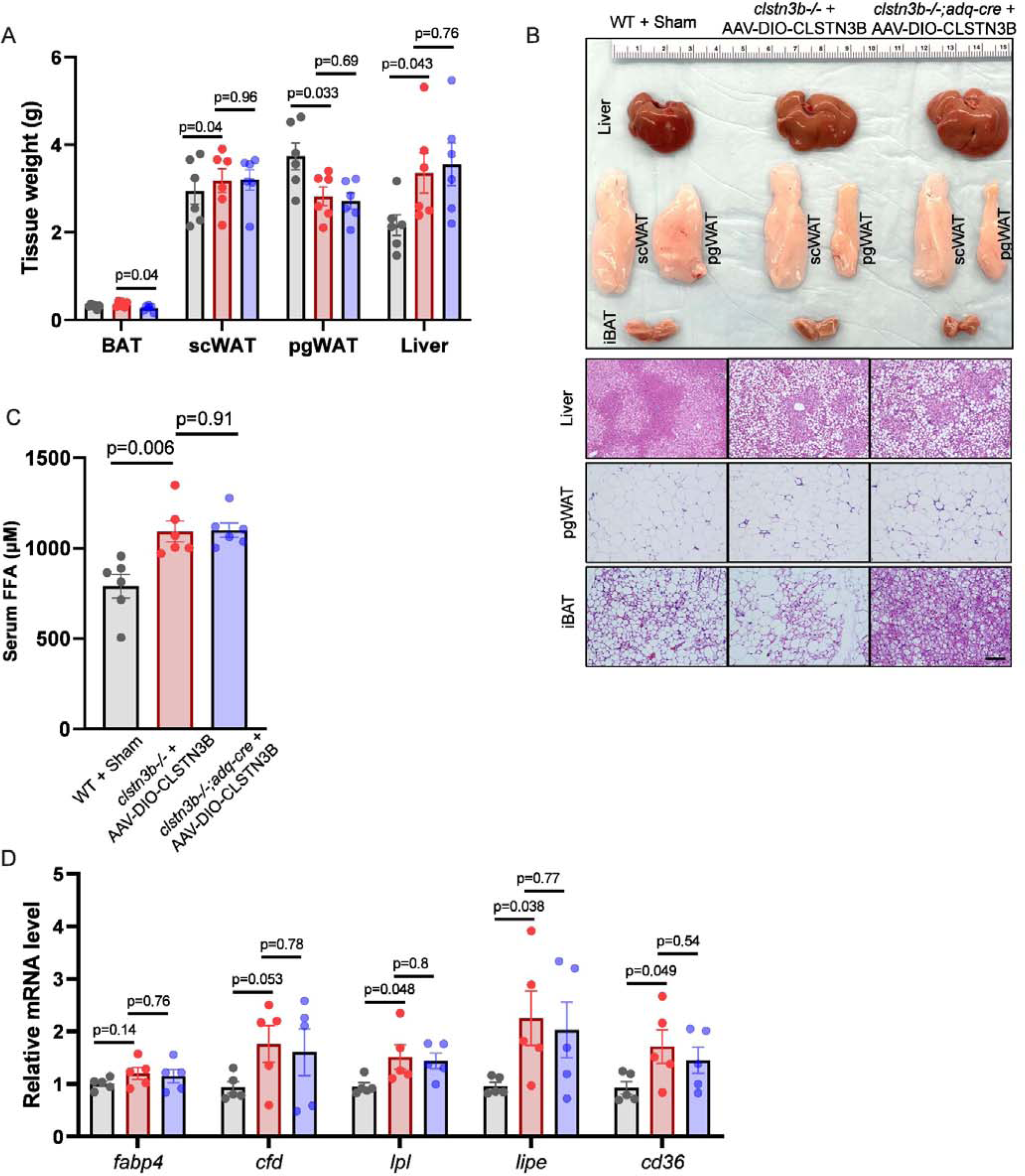
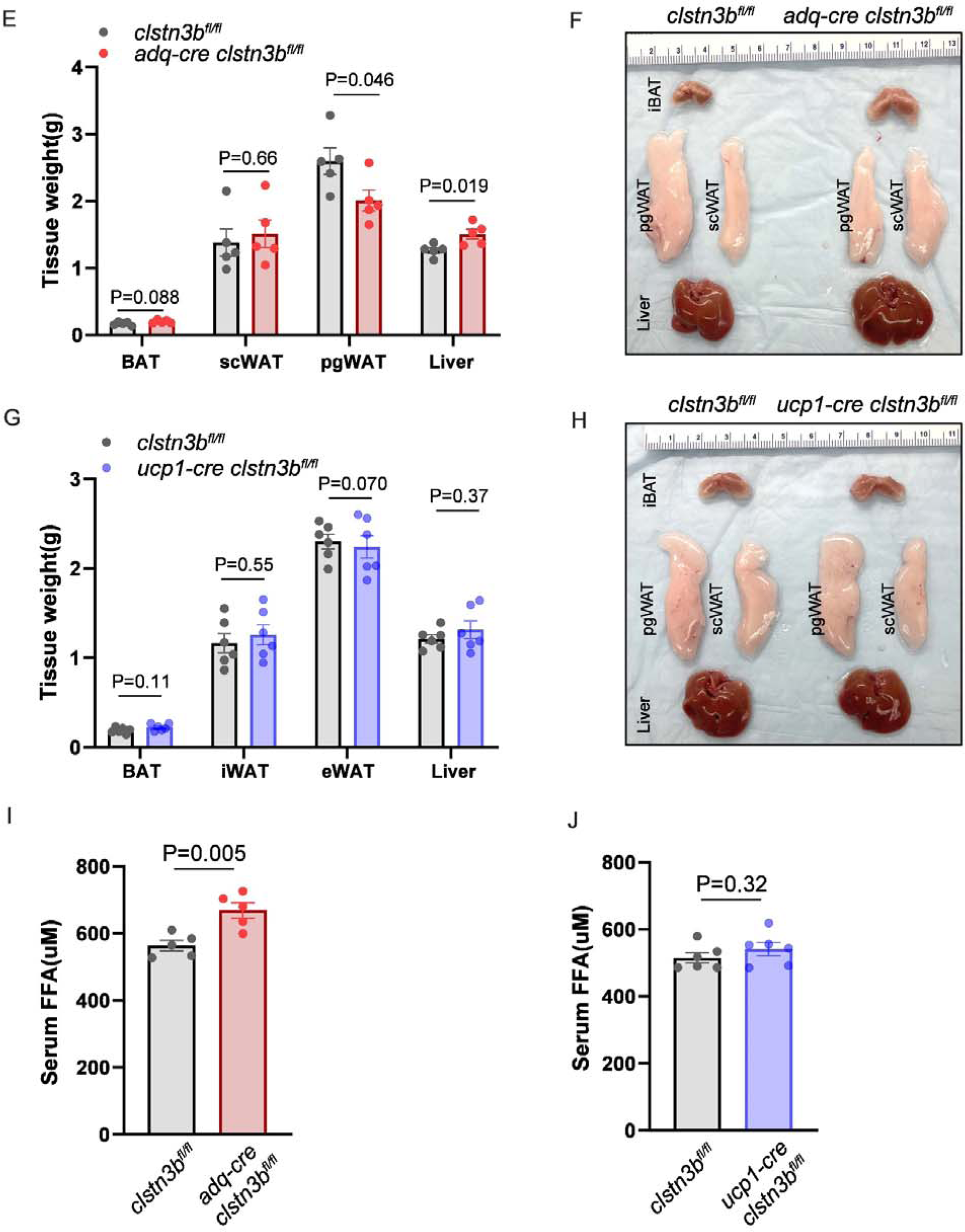
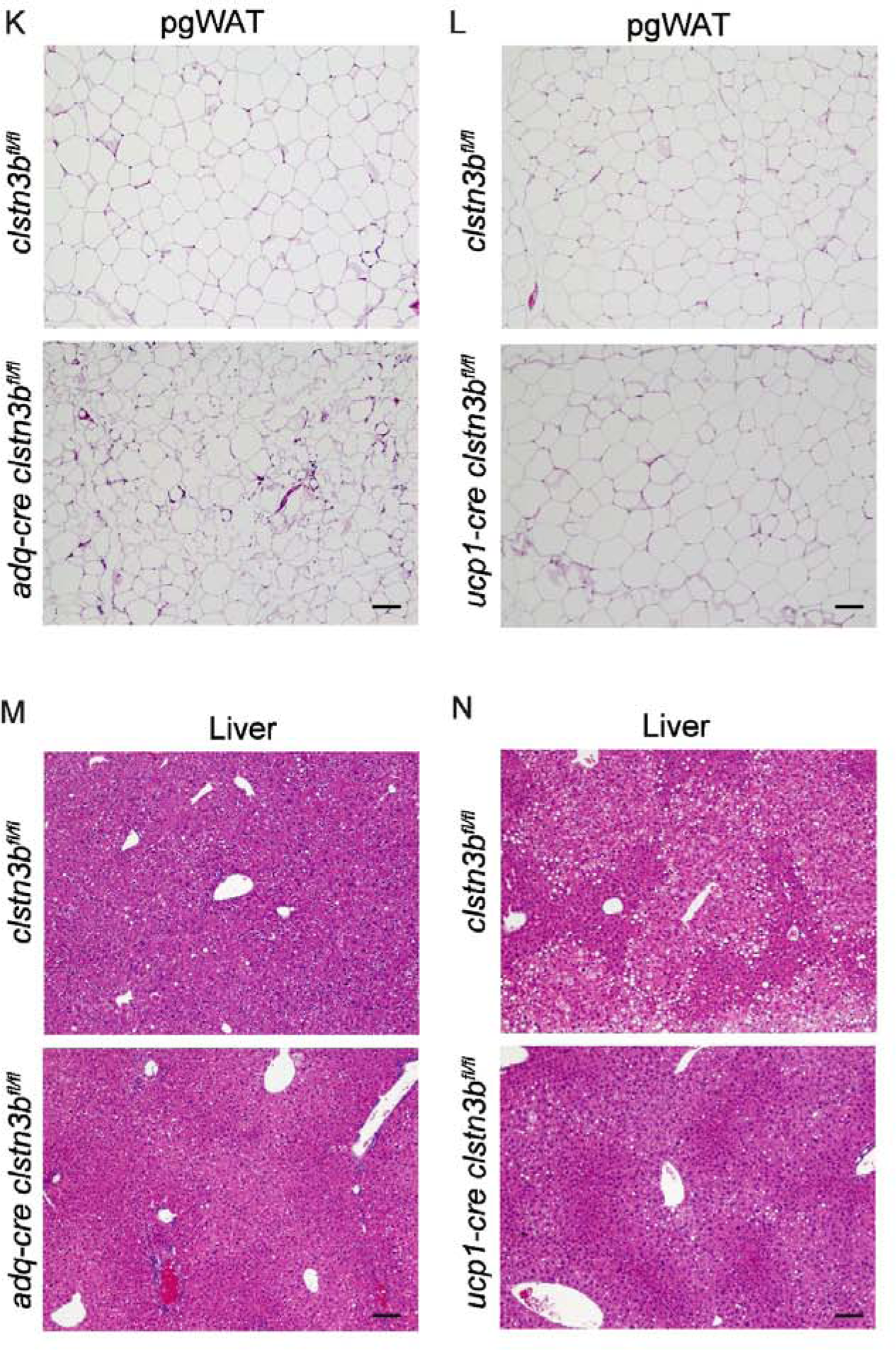
Brown adipocyte-derived CLSTN3B does not contribute to lipid storage in the WAT. A-D, Tissue weights (A), gross appearances, pgWAT, liver, and iBAT histology (B), serum FFA (C), and pgWAT gene expression analysis (D) of WT mice with sham surgery, *clstn3b-l·,* or *clstn3b-1-; adq-cre* mice receiving AAV-DIO-CLSTN3B injection into the iBAT maintained on HFD at thermoneutrality (n=6 mice). E-M, T1ssue weights (E, G), gross appearances (F, H), serum FFA (I, J), pgWAT and liver histology **(K-N)** analysis of pan-adipocyte CLSTN3B conditional KO mice *(adq-cre, clstn3bMI),* brown adipo­ cyte-specific CLSTN3B KO mice *(ucp1-cre, clstn3bfllfl)* and their ere-negative control littermates. Scale bar: 100 µm.

**Extended Data Fig 5.**
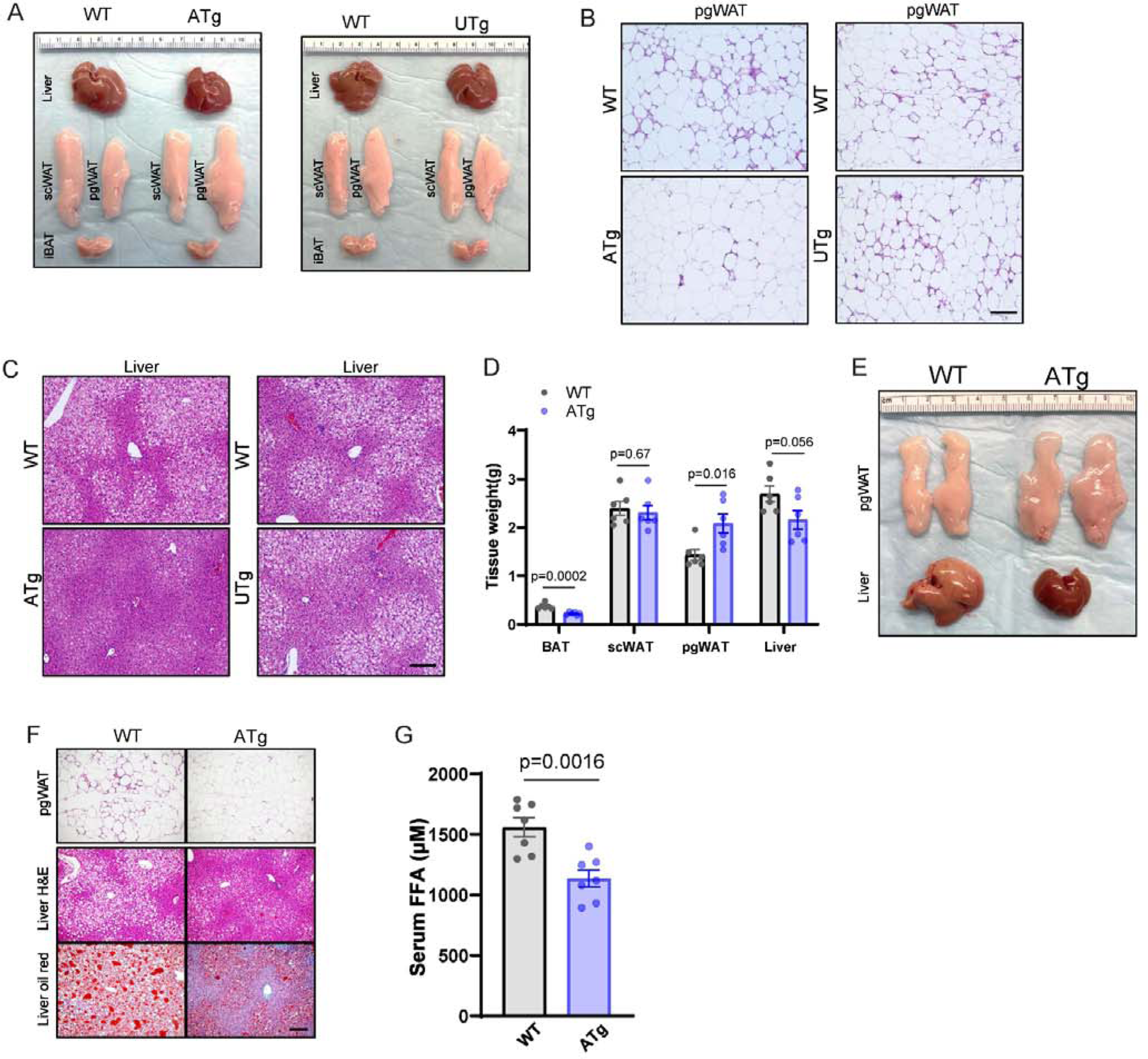
*Adq-cre* driven transgenic overexpression of CLSTN3B improves metabolic health of mice on HFD. A-C, Gross appearances (A), pgWAT (B) and liver histology (C) of *adq-cre clstn3b* transgenic mice and WT littermates on HFD at thermoneutrality. 0-G, Tissue weights (D), gross appearances **(E),** pgWAT and liver histology (F), and serum FFA (G) of *adq-cre clstn3b* transgenic mice and WT littermates on HFD at room temperature (n=6 mice).

**Extended Data Fig 6.**
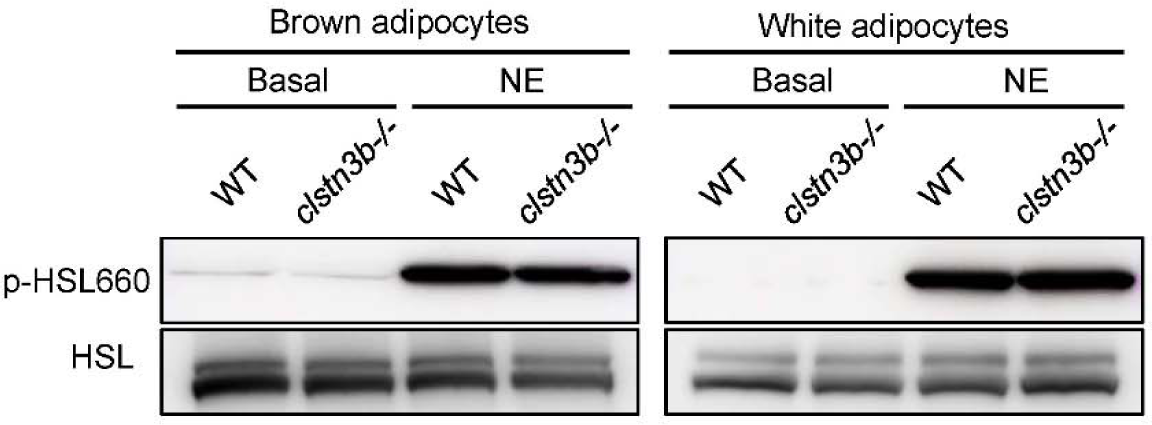
CLSTN38 ablation does not alter adipocyte HSL phosphorylation in response to NE stimulation. Western blot analysis of **HSL** phosphorylation levels in isolated *\NT* and *c/stn3b-*^1^*-* adipocytes -/+ **NE** treatment *in vitro*.

**Extended Data Fig 7.**
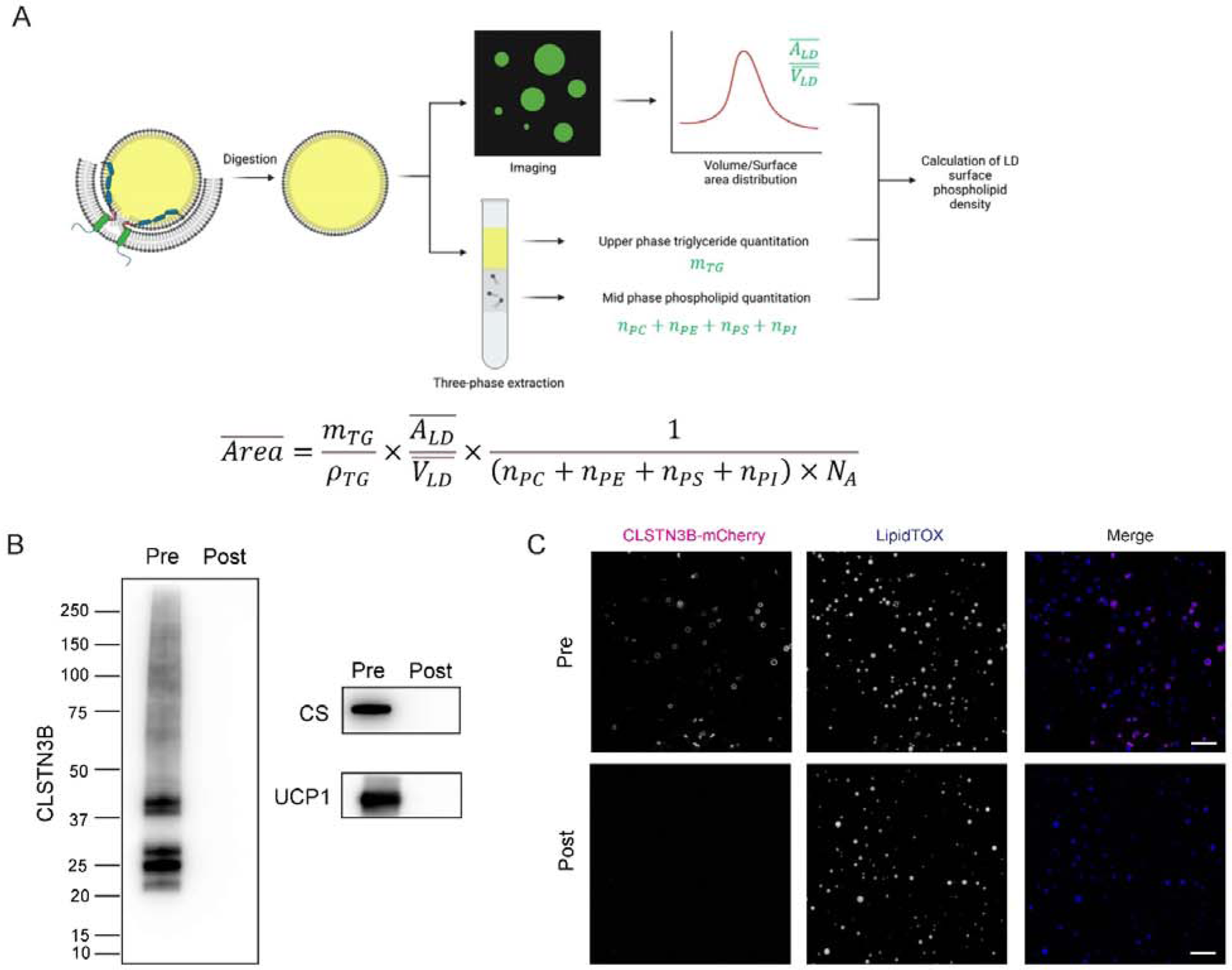
Illustration and validation of LO surface phospholipid density quantitation. A, Flowchart of LO surface phospholipid density quantitation. B, Western blot validation of removal of LO-bound ER and mitochondria by proteinase K digestion. Pre, before digestion. Note the disappearance of CLSTN3B and mitochondrial proteins in the LO fraction after proteinase K digestion. Post, after digestion. C, Fluorescence microscopy validation of removal of LO-bound ER by proteinase K digestion. Note the disap­ perance of CLSTN3B-mCherry signals around LDs after digestion. Scale bar: 5 µm.

**Extended Data Fig 8.**
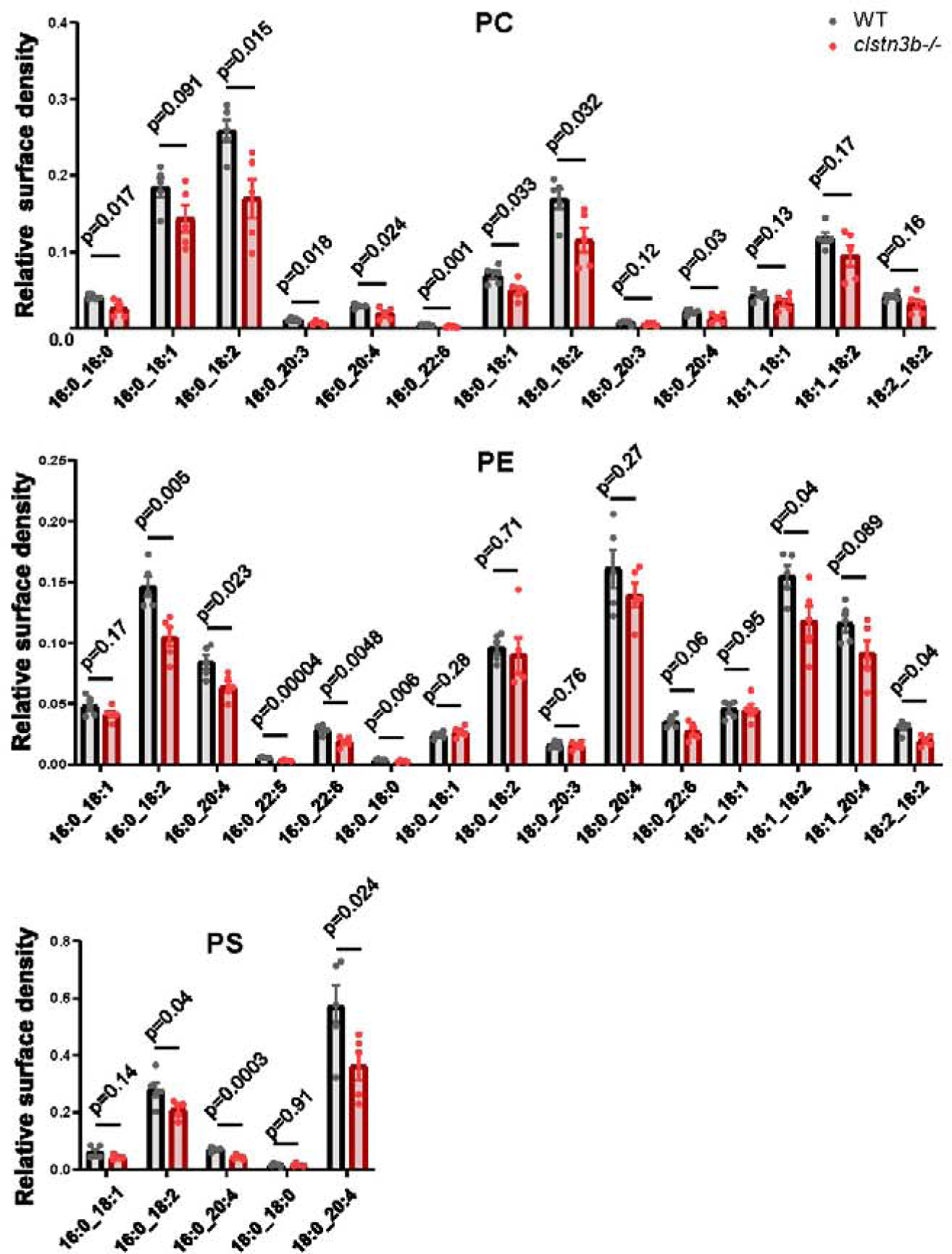
Phospholipidomics analysis of brown adipocyte LDs isolated from WT and *c/stn3b-1-* mice (n=5 replicates).

**Extended Data Fig 9.**
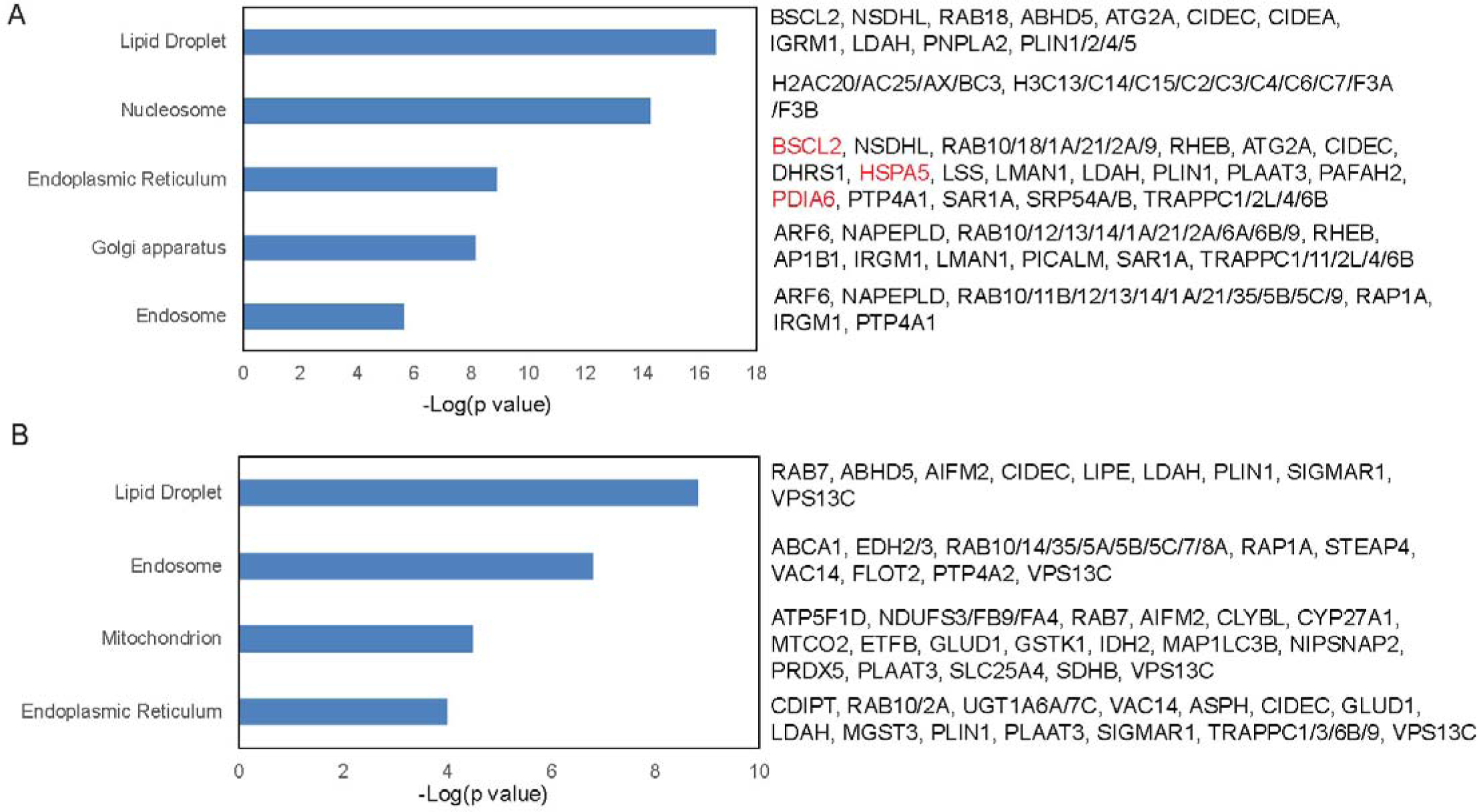
Cellular component enrichment analysis of proteins identified by proteomics analysis of adipocyte LDs isolated from WT and *clstn3b·*^1^*-* mice. A, Cellular component enrichment analysis of downregulated proteins on *clstn3b-1-* **brown adipocyte LD compared with WT. B, Cellular com­ poenent enrichment analysis of downregulated proteins on** *clstn3b-1-* **white adipocyte LD compared with WT.**

**Extended Data Fig 10.**
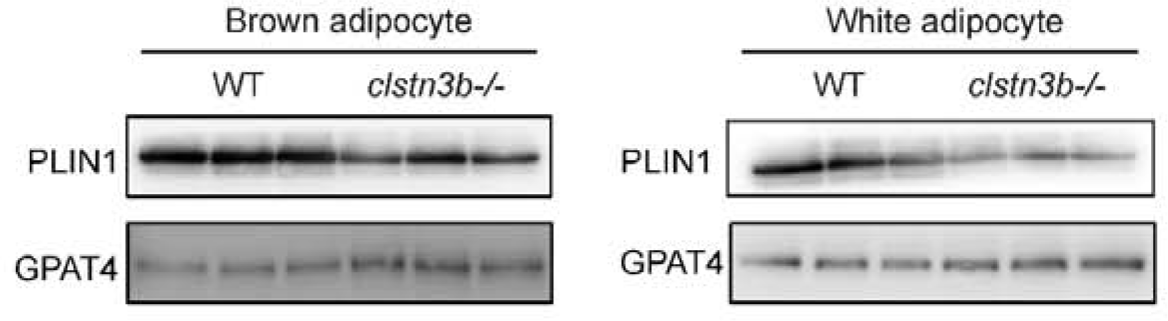
PLIN1 is downregulated on LDs isolated from *clstn3b·*^1^*•* adipocytes relative to WT adipocytes. Western blot analysis of PLIN1 protein levels in isolated LO samples from *WT* and *clstn3b-1-* adipocytes. GPAT4 was used as a loading control.

**Extended Data Fig 11.**
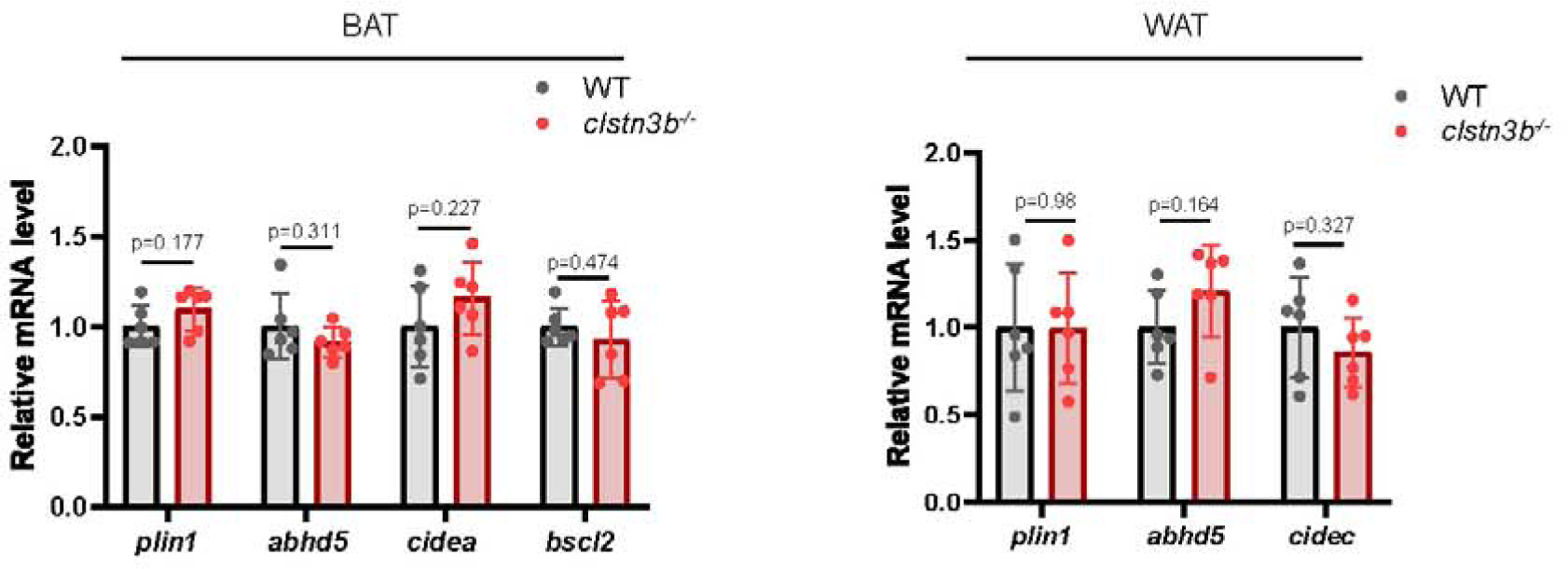
mRNA levels of downregulated LD proteins are not different between WT and *clstn3b·*^1^*•* adipose tissue. qPCR analysis of mRNA expression levels of significantly down­ regulated LD proteins in WT and *clstn3b·*^1^*-* brown and white adipose tissue.

**Extended Data Fig 12.**
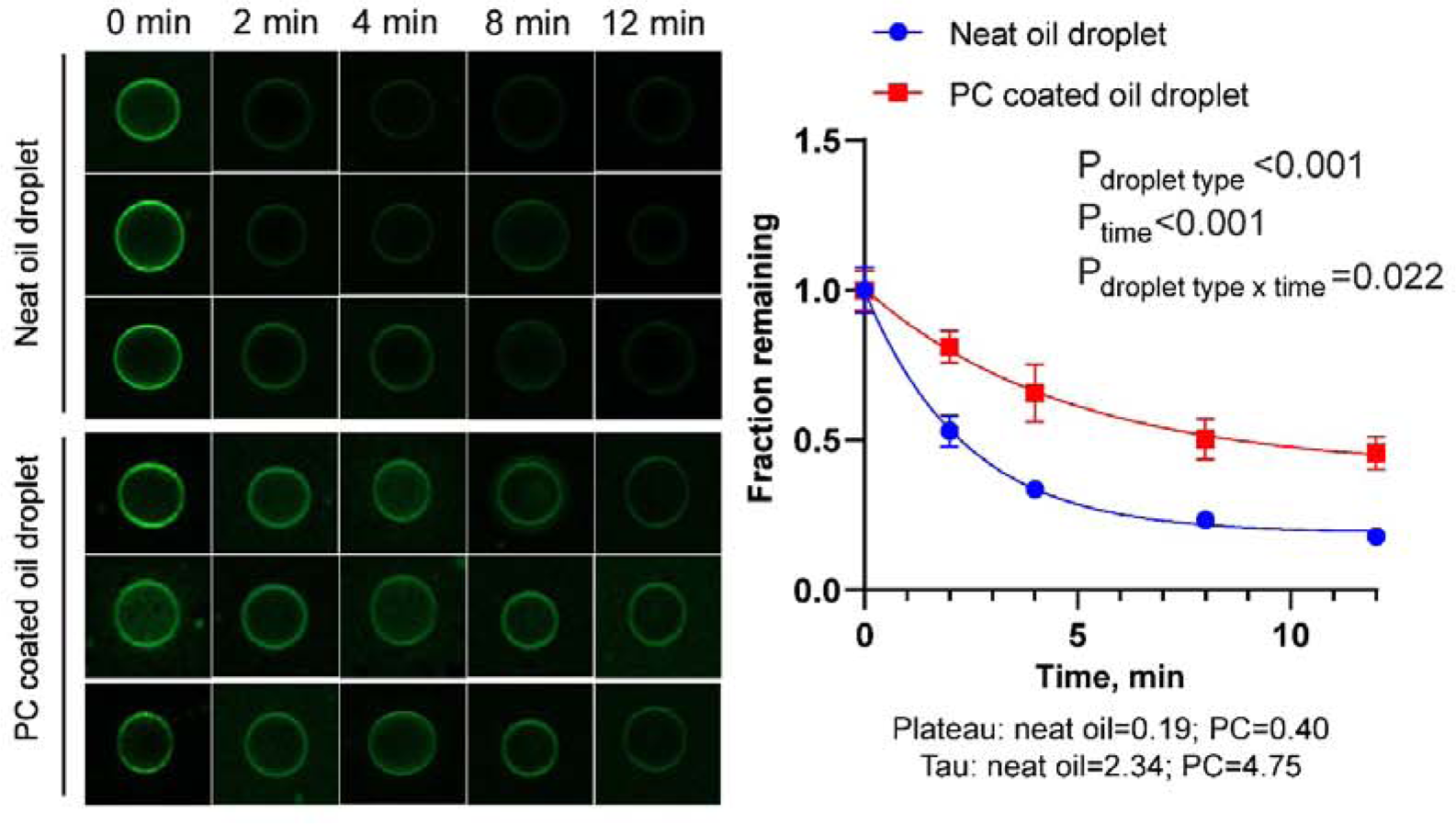
PLIN1 AH peptide dissociates more rapidly from neat triolein droplet than PC-coated triolein droplet. Trolein droplets were either emulsified with a FITC-labeled PLIN1 AH peptide or first with PC and then with the FITC-labeled PLIN1 AH peptide. The droplets were then incubated with an excess of unlabeled PLIN1 AH peptide and the fluorescence levels were recorded over time.

**Extended Data Fig 13.**
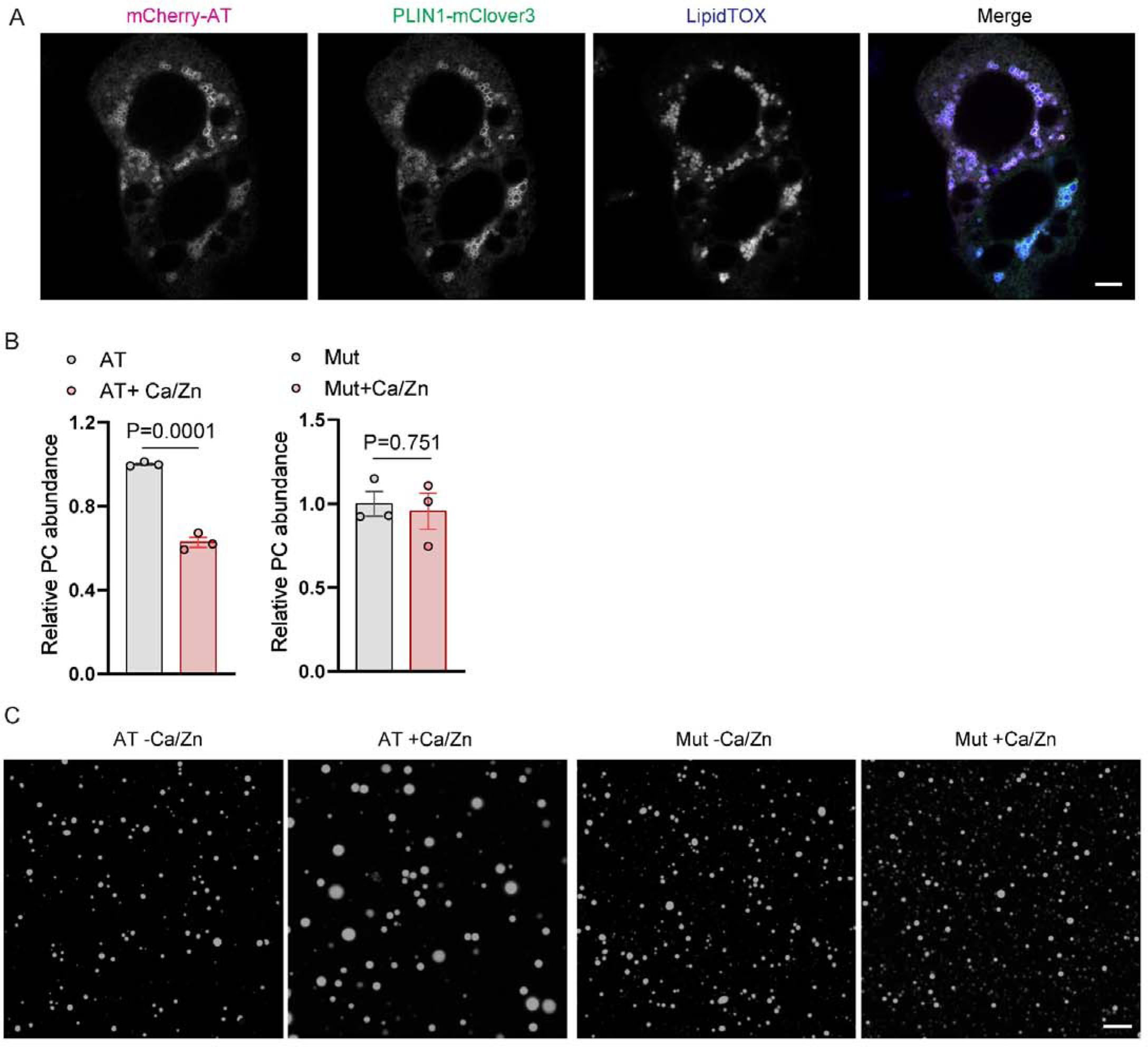
Activation of LO-targeting alpha toxin reduces LD surface phospholipid abundance. A. Fluorescence microscopy analysis of the LO-localization of CLSTN3B 1-130-mCherry-alpha toxin. Scale bar: 5 µm. B. LO phospholipid quantitation after treating isolated LDs expressing Wf or a catalytically inactive mutant alpha toxin (Mut) with or without Zn^2^+ and Ca^2^+_ Reduction in PC abundance was only observed when LDs expressing Wf alpha toxin were treated with Zn^2^+ and Ca^2^+_ C. LD size analysis after alpha toxin activation. LDs expressing Wf or mutant alpha toxin (Mut) were first treated with or without Zn^2^+ and Ca^2^+ followed by a 5-min proteinase **K** digestion. Note that significant coalescence only occurred to LDs expressing Wf alpha toxin and treated with Zn^2^• and Ca^2^+. Scale bar: 5 µm.

**Extended Data Fig 14.**
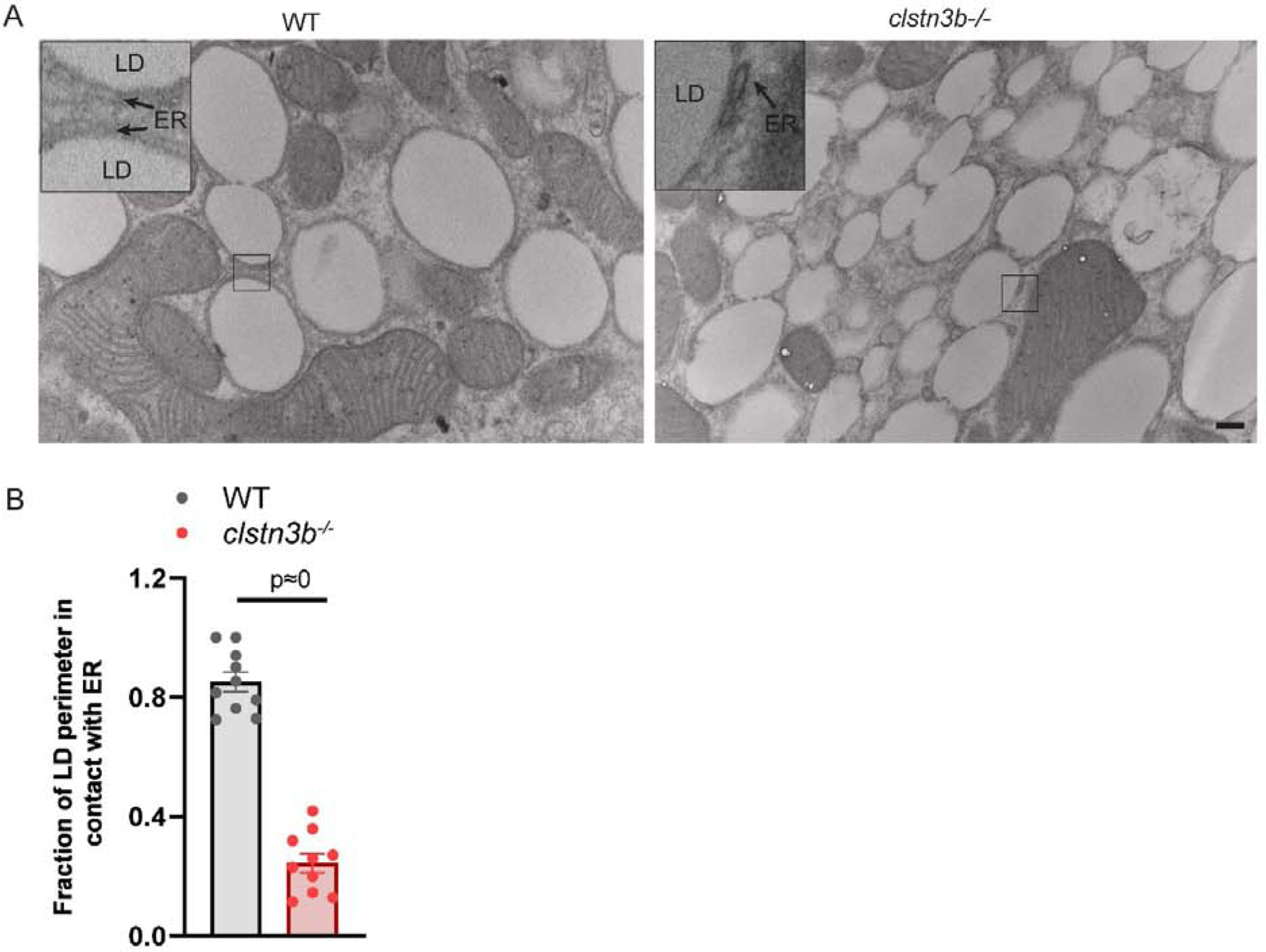
Ablation of CLSTN3B causes reduced ER association with small LDs after NE stimulation. A, Electron microscopic images of vvr and *clstn3b-l-* primary brown adipocytes showing reduced ER enwrapping of LDs in *clstn3b-l-* cells after **NE** treatment. Scale bar: 200 nm. B, Ouantitation of the extent of LO perimeter in contact with the ER in brown adipocytes (n=10 LDs).

**Extended Data Fig 15.**
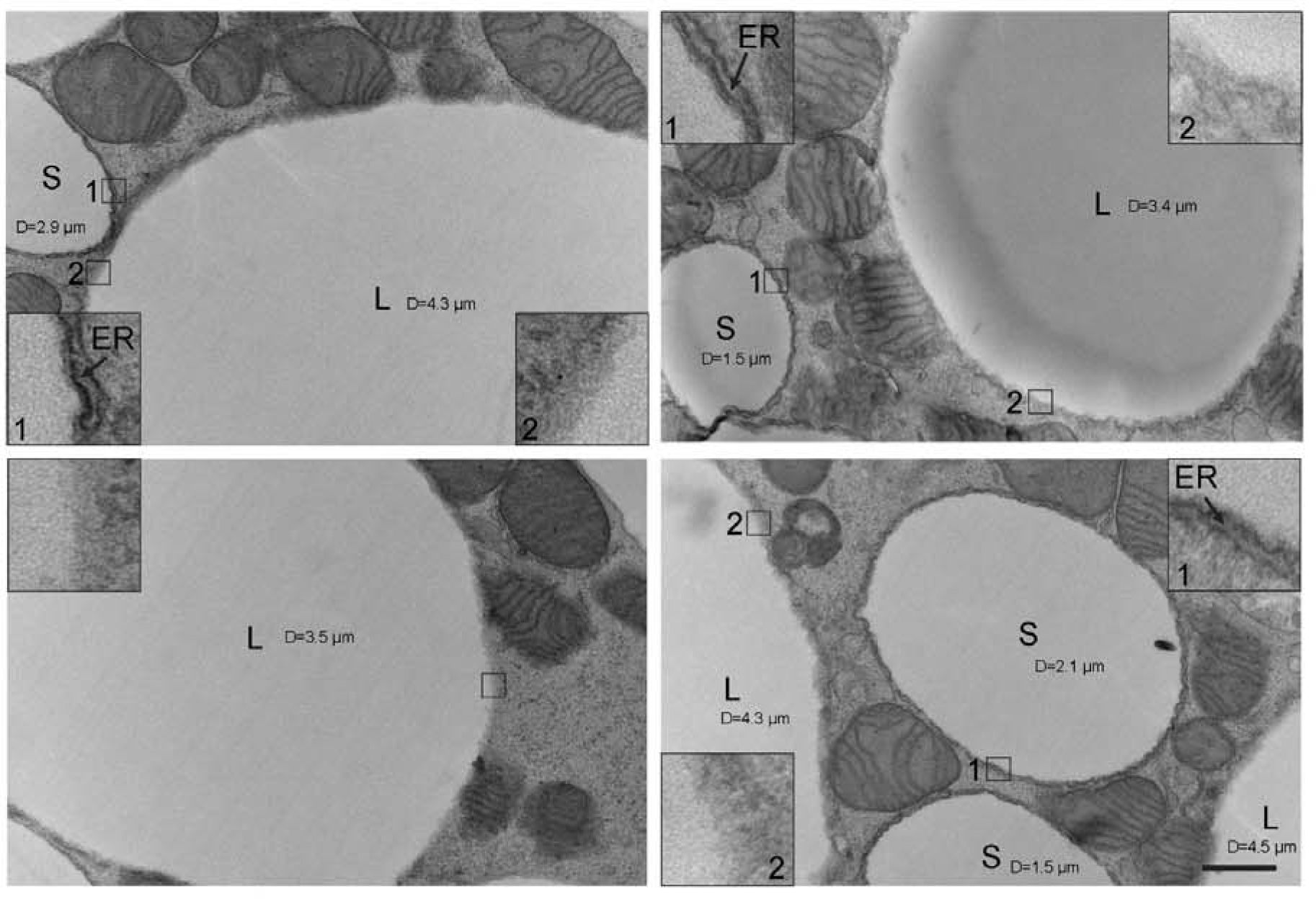
Large LDs in brown adipocytes have less extensive ER association than small LDs. Electron microscopic images of brown adipocytes showing different levels of ER association with small and large LDs. An arbitrary cutoff of D=3 µmis used to call large v.s. small LDs. S, small LD. L, large LD. Scale bar: 500 nm.

**Extended Data Fig 16.**
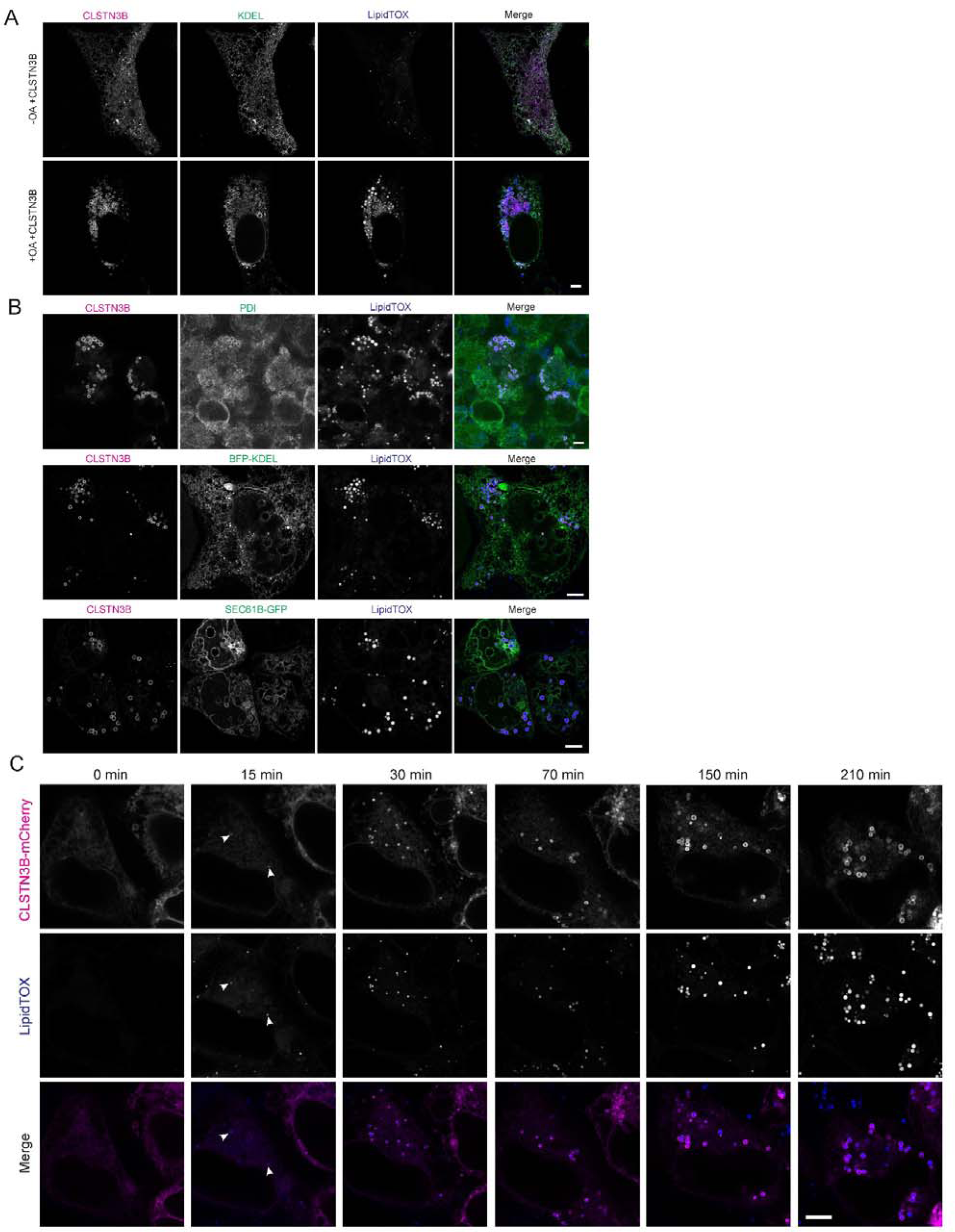
CLSTN3B promotes ER/LD contact formation in heterologous systems. A, Fluorescence microscopic analysis of CLSTN3B and KDEL localization in U2OS cells. Scale bar: 5 µm. B, Fluorescence microscopic images showing colocalization of CLSTN3B with ER markers on LO surface in HEK293 cells. Scale bar: 5 µm. C, Live imaging of HEK293 cells expressing CLSTN3B treated with OA. Imaging start­ ed immediatedly after OA addition. Note that CLSTN3B signals can be identified on nascent LDs generated within 15 min of OA addition. Scale bar: 5 µm.

**Extended Data Fig 17.**
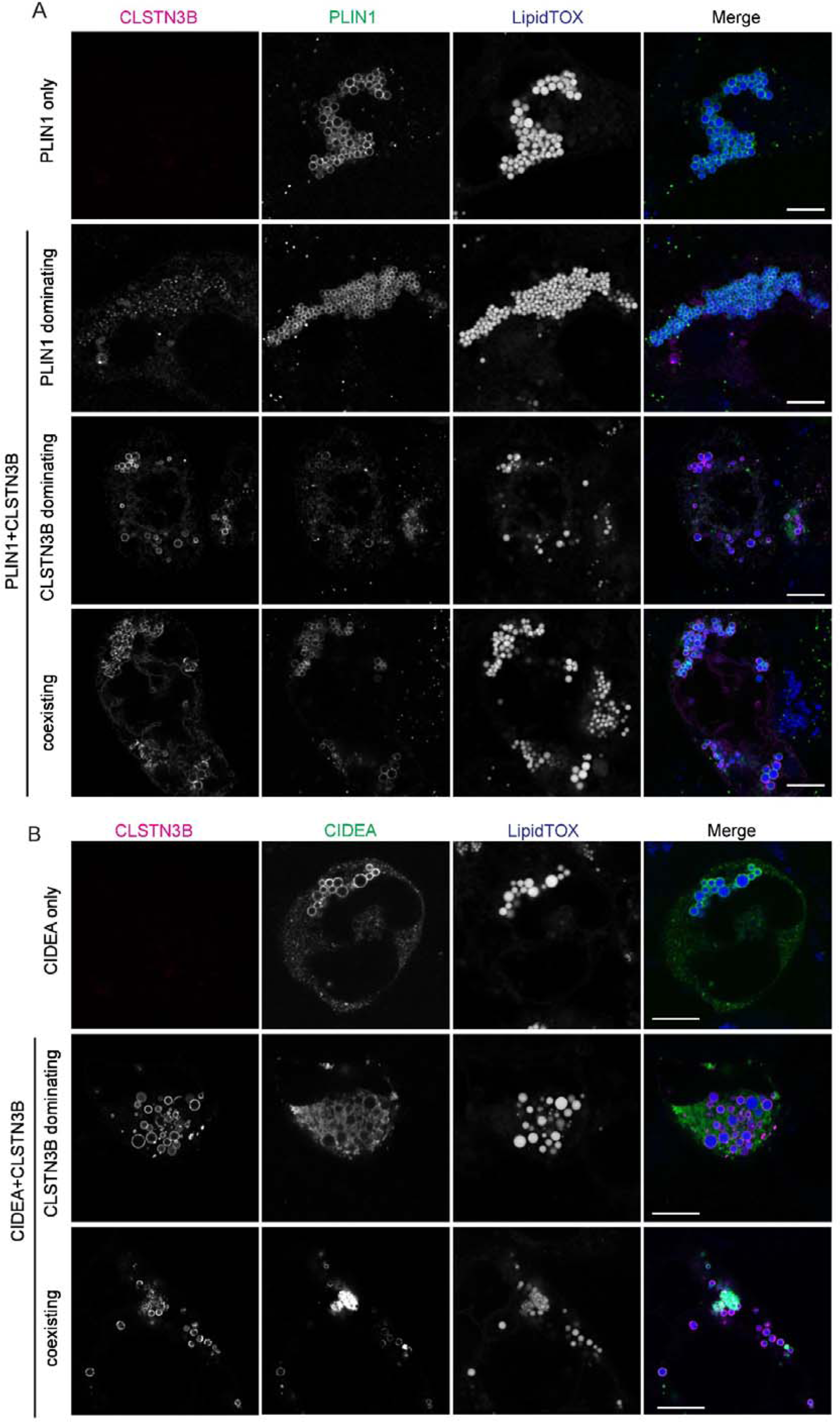
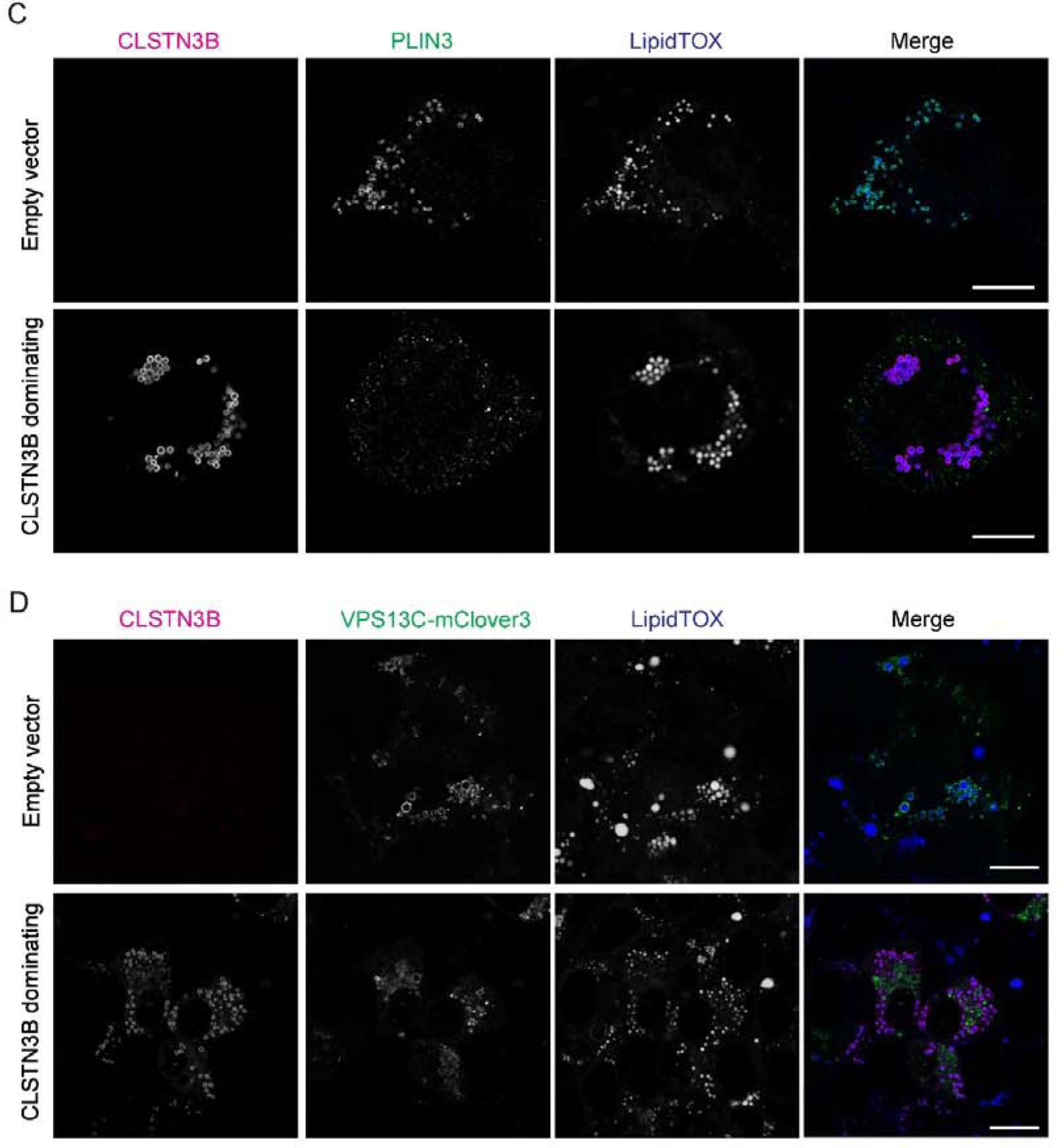
CLSTN3B displays mutually exclusive localization with AH-containing proteins on LD surface. A, Fluorescence microscopic analysis of CLSTN3B and **PLIN1** localization in HEK293 cells. PLIN1 was either expressed alone or co-expressed with CLSTN3B. Three typically observed scenarios are shown: 1) PLIN1 dominates LO localization; 2) CLSTN3B dominates LO localization; 3) PLIN1 and CLSTN3B coexist on LD surface. Complementary localization is most clearly observed under the 3rd scenario. Scale bar: 10 µm. B, Fluorescence microscopic analysis of CLSTN3B and CIOEA localization in HEK293 cells. CIOEA was either expressed alone or co-expressed with CLSTN3B. Two typically observed scenarios are shown: 1) CLSTN3B dominates LO localization; 3) CIOEA and PLIN1 coexist on LO surface. Complementary localization is most clearly observed under the 2nd scenario. Scale bar: 1O µm. C, Fluorescence microscopic analysis of CLSTN3B and PLIN3 localization in HEK293 cells. Endogenous PLIN3 was detected. Scale bar: 10 µm. 0, Fluorescence microscopic analysis of CLSTN3B and VPS13C-mClover3 localization in HEK293 cells. VPS13C-mClover3 was either expressed alone or co-expressed with CLSTN3B. Scale bar: 1O µm. CLSTN3B dominance was most commonly observed with PLIN3 and VPS13C-mClover3.

**Extended Data Fig 18.**
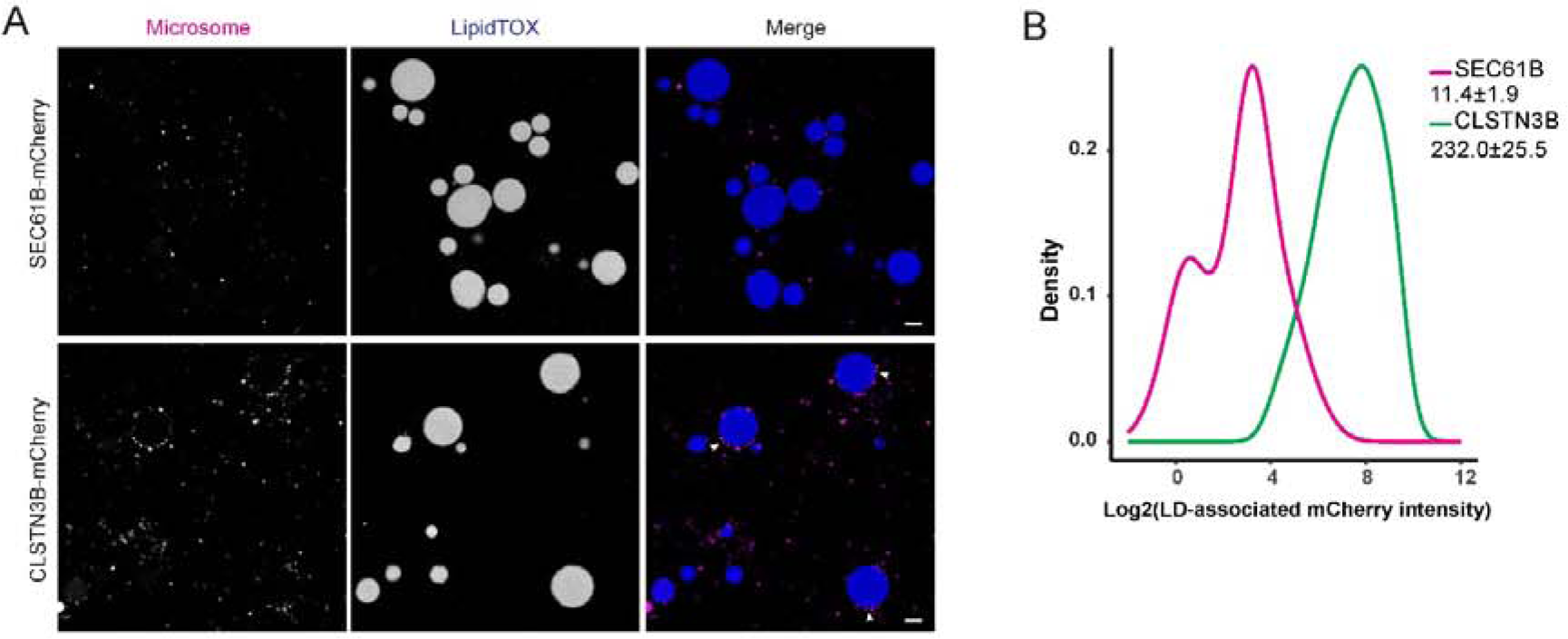
CLSTN38 promotes microsome/LD contact under a semi-reconstituted condition. A, Fluorescence microscopic analysis of microsome/LD contact. Microsomes were prepared from HEK293 cells expressing SEC618-mCherry or CLSTN38-mCherry. Scale bar: 2 µm. B, Quantitative analysis of LO-associated mCherry signals from microsomes expressing SEC618-mCherry or CLSTN3B-mCherry.

**Extended Data Fig 19.**
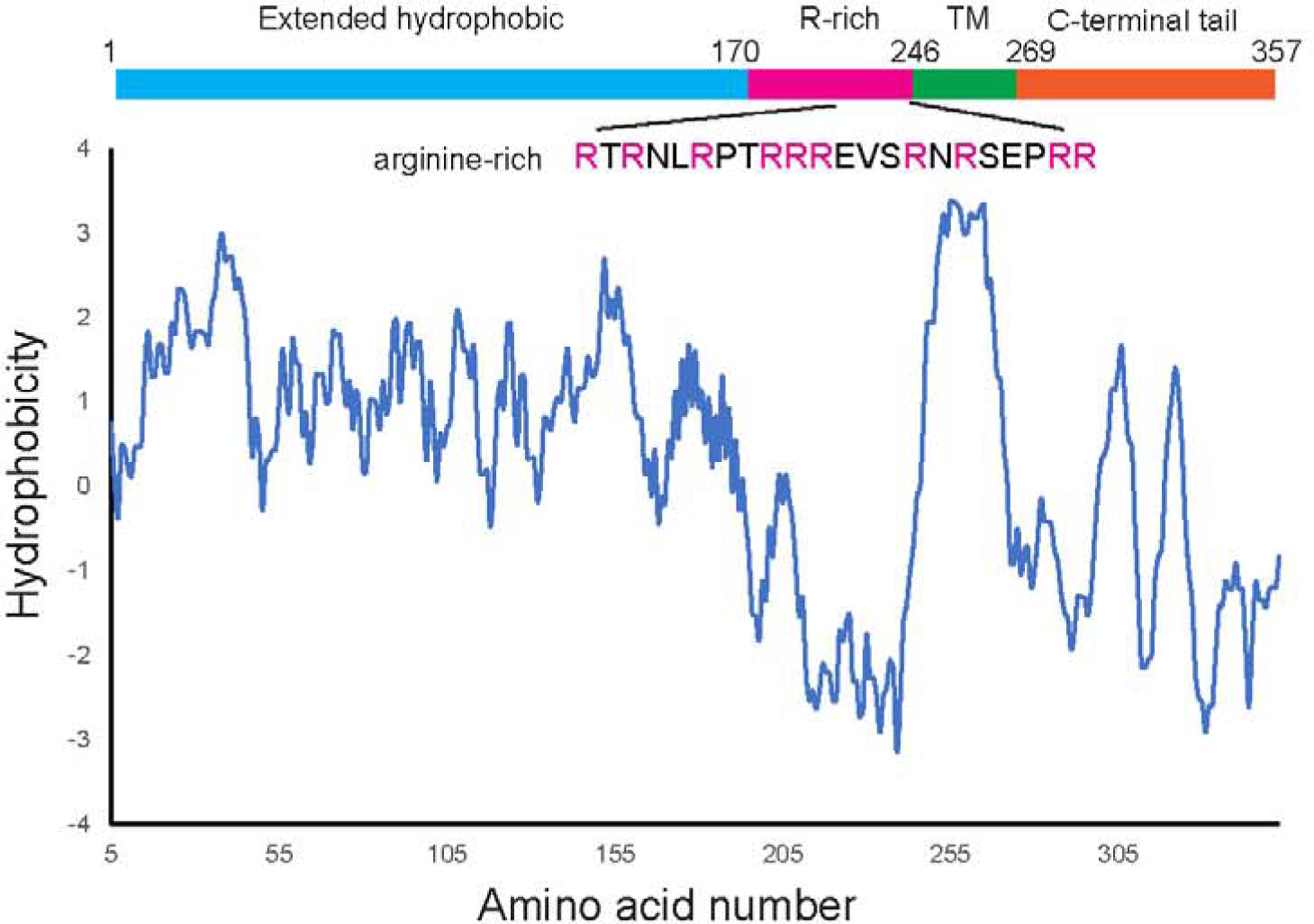
Hydrophobicity plot of CLSTN3B. The plot is generated with the Kyte and Doolittle method described in J. Mal. Biol. 157:105-132(1982).

**Extended Data Fig 20.**
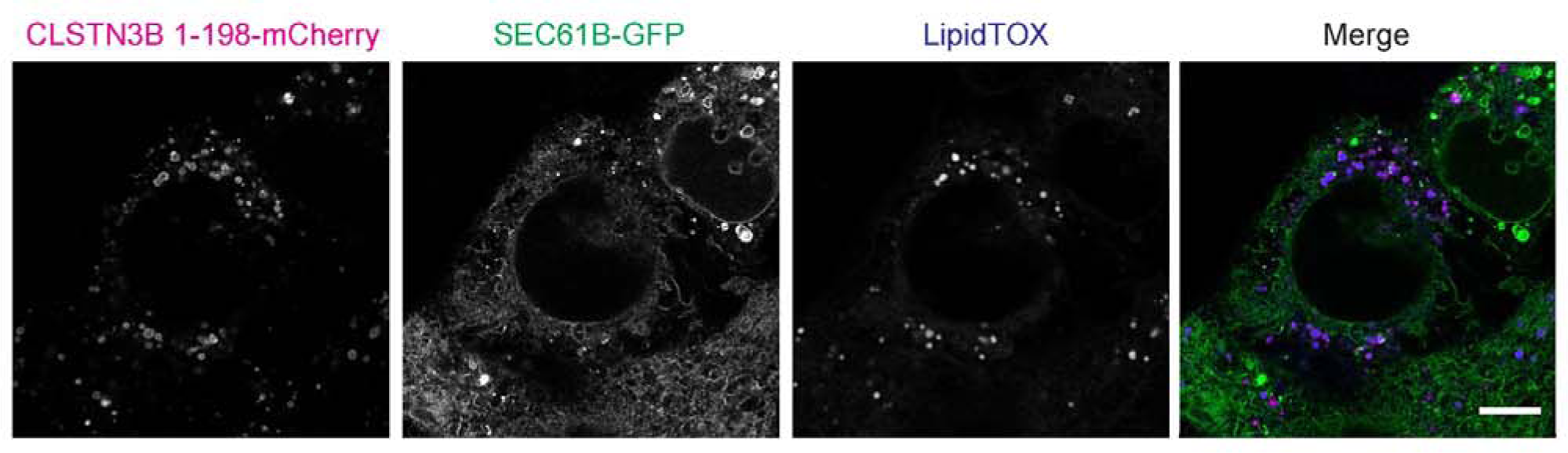
CLSTN3B 1-198 mCherry does not promote ER/LD contact formation. Fluorescence microscopic analysis CLSNT3B 1-198-mCherry and SEC618-GFP localization in HEK293 cells. Note the absence of SEC61B-GFP signal around LDs surrounded by CLSTN3B 1-198-mCherry. Scale bar: 5 µm.

**Extended Data Fig 21.**
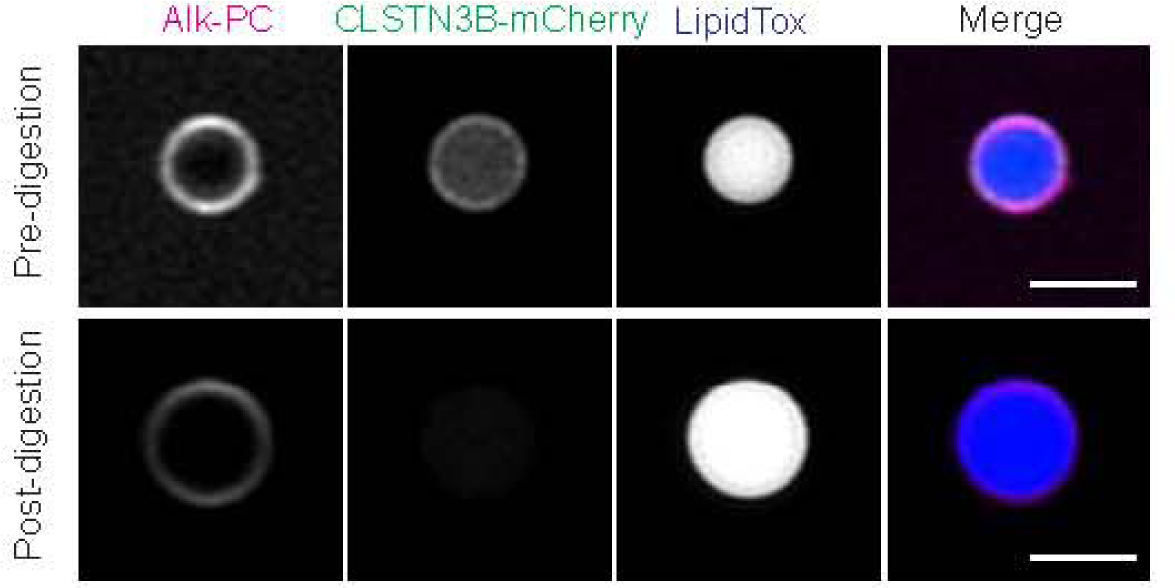
Proteinase K digestion removes LO-bound ER and allows visualization of propargyl-PC on the LD surface monolayer. LDs were harvested from HEK293 cells expressing CLSTN3B-mCherry and supplemented with propargyl-choline. Images were taken before and after proteinase K treatment. Note the absence of CLSTN3B-mCherry signal after proteinase K digestion. Scale bar: 2 µm.

**Extended Data Fig 22.**
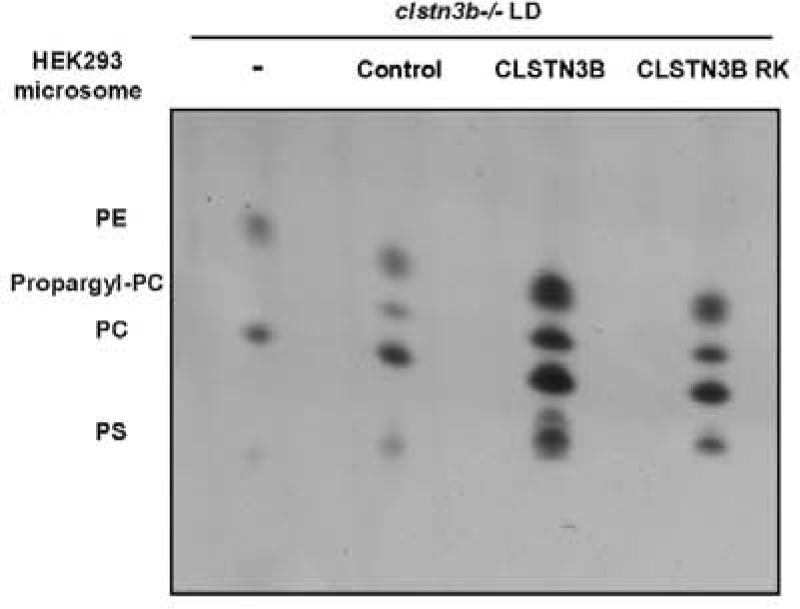
CLSTN3B promotes microsome-to-LD phospholipid transfer in a semi-re­ constituted system. TLC analysis of LO-associated propargyl-PC levels under different conditions. Note the consistency between TLC analysis and fluorescence imaging after click-chemistry labeling.

**Extended Data Fig 23.**
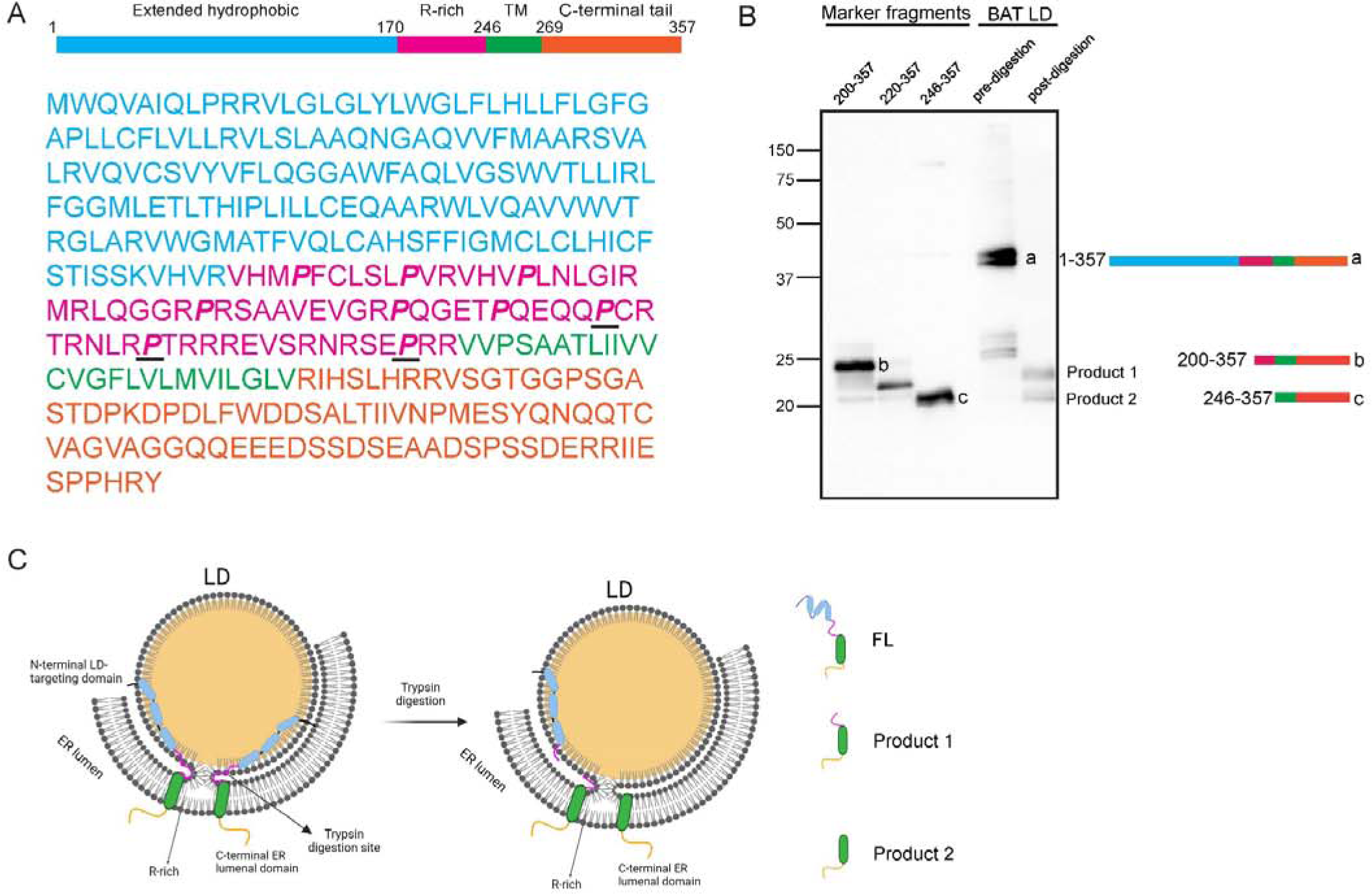
The arginine-rich region of CLSTN3B is enriched with prolines and resistant to trypsin digestion. A, Illustration of the distribution of proline residues (bold italic) in the arginine-rich region. The 3 underlined praline residues have been mutated to alanine in the P3A mutant. B, Western blot analysis of CLSTN38 trypsin digestion products from isolated brown adipocyte LD. The positions of CLSTN38 fragments with defined lengths are shown for reference. The a, b, and c landmarks are used in Fig. 6e, Schematic illustration of the BAT LD trypsin digestion results.

